# Dysregulation of *FMR1* Splicing in Human Fragile X Syndrome

**DOI:** 10.64898/2026.09.28.754997

**Authors:** Suna Jung, Joel D. Richter

## Abstract

Fragile X Syndrome (FXS) is a neuro-developmental disorder caused by a CGG expansion in *FMR1*, leading to transcriptional silencing and loss of the encoded protein FMRP. Surprisingly, ∼70% of FXS individuals express *FMR1*, but the RNA is mis-spliced to isoform *FMR1-217*, composed of exon 1 spliced to a pseudo-exon in intron 1 and cannot produce FMRP. Splice-switching ASOs rescue proper *FMR1* splicing and restore FMRP. *FMR1-217* mis-splicing increases with CGG repeat length and is negatively correlated with patient IQ. *FMR1-217* is associated with ribosome footprints, indicating it is translated into a polypeptide that may impair cognition. R-loops form at the *FMR1* locus and extend into the pseudo-exon, but splice-switching ASOs reduce *FMR1-217* and elevate FMRP independently of R-loop formation. DRB-based transcription analysis identified impaired Pol II elongation at the 5’ region of *FMR1* in FXS cells, indicated by accumulation of hypophosphorylated Pol II at the transcription start site. Consistent with this, camptothecin-induced Pol II stalling increased *FMR1-217* pseudo-exon inclusion. The splicing factors PTBP1 and PTBP2 regulate *FMR1-217* splicing in a differentiation stage-dependent manner. Together, these findings indicate that *FMR1-217* mis-splicing in FXS is associated with CGG repeat expansion, R-loop formation, impaired co-transcriptional Pol II elongation and context-dependent regulation by PTBP1/PTBP2.

## Introduction

Fragile X Syndrome (FXS) is a neurodevelopmental disorder characterized by intellectual impairment, anxiety, hyperactivity, speech and developmental delays, autism, and other co-morbidities (1). The syndrome is caused by a >200 CGG repeat expansion in the *FMR1* gene, which resides on the long (q) arm of the X-chromosome (Xq27.3). The expansion induces DNA methylation of the promoter and the expansion itself, which results in transcriptional silencing and loss of FMRP, the encoded protein. FMRP is an RNA binding protein that controls translation; its loss leads to altered protein synthesis as well as dysregulation of several downstream molecular events such as alternative splicing (2), epigenetic modifications (3), and signal transduction pathways (4,5). These and other molecular disturbances have been linked to impaired synaptic plasticity (6) and neural circuit formation (7,8), all of which are thought to underlie many of the behavioral and cognitive manifestations of the disorder.

Most investigations of FXS use *Fmr1* knockout (KO) mouse models, however, these animals do not undergo CGG expansion or DNA methylation in *Fmr1*, and thus many types of molecular dysregulation do not faithfully recapitulate what occurs in human FXS. For example, certain changes in chromatin architecture that result from the expansion, such as widespread H3K9me3 repressive marks on the X chromosome and autosomes, do not take place (9). In addition, CGG expansion-driven mis-splicing is also apparently unique to humans and possibly other higher primates. Shah et al. (10) unexpectedly found that ∼70% of FXS individuals express *FMR1* but the RNA is mis-spliced such that exon 1 is linked to a pseudo or cryptic exon within intron 1. This RNA, called *FMR1-217*, is ∼2 kb in length, is polyadenylated, and if translated could encode a 31 amino acid polypeptide. Among FXS individuals with detectable *FMR1* expression, *FMR1-217* accounts for approximately half of all *FMR1* transcripts in leukocyte samples on average. In comparison, less than 1% of *FMR1* transcripts in typically developing individuals is mis-spliced to *FMR1-217*. The importance of *FMR1-217* is suggested by the observation that it is negatively correlated with IQ. Moreover, treatment of FXS *FMR1-217*-containing cells with antisense oligonucleotides (ASO) reduce this improperly spliced form, rescue normal *FMR1* splicing, and restore FMRP. Because *FMR1-217* is also detected in FXS post-mortem brain samples, a splice-switching ASO could be a novel therapeutic approach to treat FXS.

In addition to FXS, other diseases are linked to *FMR1*, most notably Fragile X-associated Tremor/Ataxia Syndrome (FXTAS). FXTAS is a late onset neurodegenerative disorder with several co-morbidities including ataxia and impairment of executive cognition (11), which afflicts males most prominently. It develops from premutation carriers and is caused by an intermediate CGG expansion of 55-200 triplets (12). These repeats do not silence *FMR1* as in FXS; the RNA is somewhat elevated but FMRP is reduced. It has been postulated that the etiology of FXTAS is linked to translation of the CGG repeats into inclusion bodies composed of polyglycine, which may be toxic. In addition, ribosome frameshifting within the CGG expansion produces arginine-glycine peptides which may also contribute to cellular toxicity (13). A FXTAS-like CGG expansion produces *FMR1* mis-splicing to *FMR1-217* (10), indicating a molecular convergence between these two *FMR1*-related disorders.

In this study, we investigate whether *FMR1-217* might be translated and how it is generated. We demonstrate that >∼110 CGG triplets promote *FMR1-217* formation, which confirms earlier data showing that *FMR1* pre-mutation carriers display this mis-splicing event (10). The *FMR1-217* pseudo-exon is associated with ribosome footprints, indicating that the encoded 31 amino acid polypeptide is synthesized. An R-loop is formed on *FMR1* whose 3’ border extends into the *FMR1-217* pseudo-exon and increases with CGG repeat length. Although splice-switching ASOs reduce *FMR1-217* levels and restore FMRP, they do not affect R-loop formation. Assays using 5,6-dichloro-1--D-ribofuranosylbenzimidazole (DRB), which inhibits transcription elongation by abrogating Pol II serine-2 phosphorylation (14), revealed an impairment of Pol II elongation at the transcription start site of *FMR1* in FXS cells, where it resides in a hypophosphorylated state. Consistent with this, treatment of cells with camptothecin (CPT), a topoisomerase I inhibitor that induces physical stalling of elongating Pol II (15), increased *FMR1-217* mis-splicing. Sequence analysis identified several predicted polypyrimidine track binding protein (PTBP) binding sites within and flanking the *FMR1-217* pseudo-exon. Reanalysis of published CLIP-seq data confirmed PTBP2 binding within the *FMR1* CGG repeat and PTBP1/PTBP2 binding at overlapping sites in intron 1. The splicing factors PTBP1 and its paralog PTBP2 regulate *FMR1-217* splicing in a differentiation stage-dependent manner. Our data indicate that *FMR1* mis-splicing to generate *FMR1-217* is regulated by multiple converging mechanisms including CGG repeat expansion, R-loop formation, perturbation of co-transcriptional Pol II elongation, and context-dependent regulation by PTBP1/PTBP2. These data suggest that dysregulated RNA processing at the *FMR1* locus generates a disease-relevant isoform that may contribute to the cognitive impairment of FXS.

## Materials and Methods

### Cell culture

EBV-transformed human lymphoblastoid cells (LCLs) were obtained from Coriell Institute. Two control lines from typically developing (TD) individuals, GM07174 (Con1) and GM06890 (Con2) and two premutation carrier patient-derived lines, GM20233 (repeat size of 117), GM06891 (repeat size of 118); and two FXS patient-derived lines, GM07365 (FXS1), GM06897 (FXS2) were used in this study. LCLs were cultured in RPMI 1640 medium (Gibco, Cat # 11875127) supplemented with 15% FBS (Gibco, Cat # A5256701) and 2 mM L-glutamine (Gibco, Cat # 25030081) and 1% antibiotic-antimycotic solution. Cells were maintained at a density of 2 x 10^5^ viable cells/ml in T25 flasks. HEK293T cells were cultured in DMEM (Gibco, #11995065) containing 10% FBS and 1% antibiotic-antimycotic solution.

Control human foreskin fibroblasts (HFFs) were obtained from Dr. Megan Orzalli (University of Massachusetts Chan Medical School). FXTAS and FXS patient-derived fibroblasts were obtained from Dr. Elizabeth Berry-Kravis (Rush University). FXTAS fibroblast line F35, which is mosaic and carries 32, 121, and 135 repeats, and line C172 carries 140 CGG repeats. FXS fibroblasts harbor >200 CGG repeats and exhibit 81% *FMR1* methylation. Fibroblasts were cultured in DMEM medium supplemented with 10% FBS and 1% antibiotic-antimycotic solution.

Typically developing human iPSCs were obtained from Dr. Sandra Almeida (University of Massachusetts Chan Medical School) and TC43-97 unmethylated full mutation FXS iPSCs carrying approximately 270 CGGs were obtained from Dr. Peter Todd (University of Michigan) as previously described in Rodriguez et al. (16). iPSCs were maintained on Matrigel (Corning® Matrigel® hESC-Qualified Matrix, Cat # 354277)-coated six well plates in mTesR Plus media (STEMCELL Technologies, Cat # 100-0276) and passaged using Gentle Cell Dissociation Reagent (GCDR, STEMCELL Technologies, Cat # 100-1077). 10 μM Y-27632 ROCK inhibitor (EMD Millipore, Cat # 688001) was added to the culture medium when passage the cells.

Human iPSCs were differentiated into neural progenitor cells (NPCs) using a monolayer-based neural induction method. Briefly, iPSCs were dissociated into single cells using Accutase (STEMCELL Technologies, Cat # 07920) and seeded at 2 x 10^6^ cells per well of a Matrigel coated 6-well plate in STEMdiff^TM^ Neural Induction Medium (STMECELL Technologies, Cat # 05839) supplemented with 10 μM Y-27632 ROCK inhibitor for the first 24 hours. Cells were cultured for 3 weeks with passaging performed once per week using Accutase. Following neural induction, NPCs were expanded and maintained in STEMdiff^TM^ Neural Progenitor Medium (STEMCELL Technologies, Cat # 05833).

For neuronal differentiation, NPCs were plated onto poly-L-ornithine (PLO)/laminin-coated plates and differentiated into forebrain neurons using the STEMdiff^TM^ Forebrain Neuron Differentiation Kit (STEMCELL Technologies, Cat #0 8600) and Maturation Kit (STEMCELL Technologies, Cat # 08605), according to the manufacturer’s protocol. Cells were differentiated for 7 days, followed by 8 days of maturation.

### Immunofluorescence staining of NPCs

NPCs were fixed with 4% paraformaldehyde (PFA), followed by three washes with PBS. Cells were permeabilized with 0.1% Triton X-100, washed three times with PBS, and blocked for 1 h at RT in 3% BSA in PBS. Cells were incubated overnight at 4 °C with primary antibody against Nestin (STEMCELL Technologies, Cat # 60091, 1:250), followed by three washes with PBS. Cells were subsequently incubated with goat anti-mouse Alexa Fluor^TM^ 488 secondary antibody (Thermo Fisher Scientific, Cat # A28175, 1:300) for 1h at RT, washed three times with PBS, and nuclei were counterstained with Hoechst 33342 (Thermo Fisher Scientific, Cat # H3570). Images were acquired using a Leica SP8 confocal microscope.

### siRNA transfection

Cells were transfected with siRNA against PTBP1 (IDT, Desgin ID # hs.Ri.PTBP1.13.2), PTBP2 (IDT, Design ID # hs.Ri.PTBP2.13.2), TARDBP (Dharmacon, Cat # L-012394-00-0005), SFPQ (IDT, Design ID # hs.Ri.SFPQ.13.1) or non-targeting control siRNA (IDT, Negative Control DsiRNA, Cat # 51-01-14-04) using Lipofectamine RNAiMAX (Thermo Fisher Scientific, Cat # 13778075) for 72 h following the manufacturer’s instructions. For siRNA transfection to iPSCs, cells were transfected as previously described in Ma et al. (17). ROCK inhibitor was added 1h prior to single cell dissociation. Cells were dissociated into single cells using GCDR, and 4 x 10^5^ viable cells were resuspended in a transfection solution containing 30 pmol siRNA and Lipofectamine RNAiMAX. The cell suspension was incubated for 10 min at room temperature and then seeded onto Matrigel-coated 12 well plates in fresh mTesR Plus medium supplemented with ROCK inhibitor.

### FMR1-217-FLAG cloning and overexpression

FMR1-217 DNA was amplified from FXS LCLs using Q5 High-Fidelity 2X Master Mix (NEB, Cat # M0492) with cloning primers 217F and 217R, using the following PCR conditions: initial denaturation at 98°C for 2 min; 35 cycles of 98°C for 10 s and 72°C for 1 min; and a final extension at 72°C for 2 min. PCR products were resolved by gel electrophoresis, and the target band was excised and purified using the Monarch® DNA Gel Extraction Kit (NEB, Cat # T1020S). A 3xFLAG tag flanked by restriction enzyme sites was synthesized as the reverse primer 3XFLAG-R (Oligo-Flex Services, Genewiz), and fused to the 3’ end of FMR1-217 by PCR using primers 217F and 3xFLAG-R. The PCR product was resolved on a 2.5% agarose gel, excised, and purified using the QIAquick PCR Purification Kit (Qiagen, Cat # 28106). The resulting FMR1-217-3xFLAG insert and pCAG-tdTomato backbone (NEB, Cat # 432868) were both digested with EcoRI and BglII, and ligated using NEB Quick Ligase (NEB, M2200S). The ligation product was transformed into NEB Stable competent E. coli.

Plasmids were purified using the Plasmid Plus Midi Kit (Qiagen, Cat # 12943). HEK293T cells were transfected with 2 μg of pCAG-tdTomato or pCAG-FMR1-217-3xFLAG using Lipofectamine 3000 (Thermo Fisher Scientific, Cat # L3000015). Cells were harvested 48 hours post-transfection, and FLAG-tagged petide expression was assessed by Western blot.

### ASO treatment

For ASO treatments, Con1 and FXS1 LCLs were seeded at a density of 6 x 10^5^ viable cells in 3 ml of culture medium in the T25 flasks. On day 1, cells were transfected with 80 nM of each ASO (713 and 714) using Lipofectamine RNAiMAX (see Shah et al. (10) for details of the ASOs). On days 2 and 5, 1 μM 5-AzadC (Millipore Sigma, Cat # A3656) was added to the culture medium. Cells were harvested on day 7 for downstream analysis.

### Transcription elongation dynamics assay

For DRB washout experiments, LCLs were treated with 100 μM DRB (Millipore Sigma, Cat # 287891) or DMSO (Millipore Sigma, Cat # D2650) for 3 h and collected by centrifugation at 300 x g for 5 min. The cell pellets were washed with cold PBS and resuspended in pre-warmed fresh RPMI 1640 medium to initiate transcriptional recovery. Cells were harvested 5 min intervals to monitor transcription accumulation across the *FMR1* locus. To assess the effect of Pol II stalling on *FMR1* splicing, LCLs were treated with 10 μM CPT or DMSO for 1 h before harvesting. Total RNA was extracted and transcript levels were quantified by qPCR.

### RNA extraction, cDNA synthesis, qPCR

Cells were washed with PBS and collected by centrifugation. Total RNA was extracted using TRIzol reagent (Invitrogen, Cat # 15596018) according to manufacturer’s instructions. Briefly, TRIzol was added to lyse the cells and incubated for 3 min at room temperature. Chloroform was added and samples were vortexed before centrifugation at 13,000 rpm for 10 min at 4 °C. The aqueous phase was collected, and RNA was precipitated overnight with isopropanol in the presence of glycogen (Thermo Scientific, Cat # AM9510). RNA was centrifugated at 13,000 rpm for 30 min at 4 °C, washed twice with 70% ethanol, and resuspended in RNase-free water (Invitrogen, Cat # 1097702). For cDNA synthesis, 1 μg of total RNA was reverse transcribed using QuantiTect Reverse Transcription Kit (Qiagen, Cat # 205311) following the manufacturer’s protocol. Quantitative PCR (qPCR) was performed using SYBR green Master Mix (Bio-Rad, Cat # 1725122) on a QuantStudio3 Real-Time PCR System under following conditions: 50 °C for 2 min, 95 °C for 10 min, followed by 39 cycles of 95 °C for 15 s, 60 °C for 60 s.

### Primers

The sequences of all primers used in this study, including sequences from prior work (18–22), are provided in Supplementary Table S1 and S2.

### Western blot analysis

Cells were lysed in RIPA buffer supplemented with protease inhibitor (cOmplete^TM^ EDTA-free, Sigma) and phosphatase inhibitor cocktails (PhosSTOP^TM^, Roche, Cat # 4906845001). Lysates were rotated at 4 °C and centrifugated at 13,000 rpm for 10 min at 4 °C. The supernatants were collected, and protein concentration was measured using Pierce BCA Protein Assay Kit (Thermo Fisher Scientific, Cat # 23225). Protein samples were mixed with 4x Laemmli buffer (Bio-Rad, Cat # 1610747) and denatured at 95 °C for 5 min. Samples were separated on 10% SDS PAGE gels and transferred onto PVDF membranes by using a Trans-Blot Turbo semi-dry transfer system (Bio-Rad). Membranes were blocked in blocking solution (Bio-Rad) for 1h at room temperature and washed three times for 10 min with PBST. Membranes were incubated overnight at 4 °C with the primary antibodies: FMRP (Millipore, Cat # MAB2160), GAPDH (Cell Signaling Technology, Cat # 2118), alpha-Tubulin (Sigma, Cat # T5168), p-Ser5 Pol II (Abcam, Cat # ab5408), PTBP1 (Cell Signaling Technology, Cat # 57246), PTBP2 (Cell Signaling Technology, Cat # 59753), TDP-43 (Proteintech, Cat # 10782-2-AP). After washing three times with PBST, membranes were incubated with HRP conjugated anti-mouse (Thermo Fisher Scientific, Cat # 31430) or anti-rabbit (Thermo Fisher Scientific, Cat # 31460) secondary antibodies. Protein bands were detected using enhanced chemiluminescence (ECL, Revvity Health Sciences Inc, Cat # 50-904-9326) reagents and visualized with an Amersham imaging system.

### DNA-RNA immunoprecipitation (DRIP)-qPCR

DRIP-qPCR was performed as described in Sanz et al. (18). Approximately 6 x 10^6^ cells were used when available. Cells were lysed in TE buffer containing 20% SDS and proteinase K (Roche, Cat # 3115828001) and incubated overnight at 37 °C. Genomic DNA was extracted using phenol/chloroform/isoamyl alcohol (25:24:1) (Invitrogen, Cat # 15593031) and precipitated with sodium acetate (NaOAc) and ethanol. DNA was spooled, washed with 80% ethanol and resuspended in TE buffer. Genomic DNA was then digested with restriction enzyme cocktails of EcoRI (NEB, Cat # R3101S), HindIII (NEB, Cat # R3104T), BsrGI (NEB, Cat # R3101S), and XbaI (NEB, Cat # R0145S) in the presence of spermidine at 37 °C overnight. Fragmented DNA was extracted using phase-lock gel tubes with phenol/chloroform/isoamyl alcohol (25:24:1) at 16,000g for 10 min at 4 °C. The aqueous phase was collected, and DNA was precipitated with NaOAc and ethanol for 1h at least, followed by centrifugation at 16,000g for 35 min at 4 °C. The resulting DNA pellet was washed with 80% ethanol and resuspended in TE buffer. As a control, RNaseH (NEB, Cat # M0297L) was treated for overnight at 37 °C.

For immunoprecipitation, 4 μg of digested DNA was incubated with 4 μg S9.6 antibody (Kerafast, Cat # ENH001) in DRIP binding buffer at 4 °C overnight. 25 μl of each Dynabeads Protein A (Invitrogen, Cat # 10001D) and G (Invitrogen, Cat # 10003D) was added and rotated at 4 °C for 2h. Beads bound to DNA were washed three times in DRIP wash buffer by rotating 10 min at room temperature. After the final wash, DRIP elution buffer and proteinase K were added and samples were incubated at 55 °C for 45 min in thermomixer at 300 rpm. Beads were then removed using magnetic stand for 3 min. DNA was extracted with phenol/chloroform/isoamyl alcohol (25:24:1), centrifugated, and pellets were air-dried and resuspended in 50 μl of RNase free water.

### Chromatin immunoprecipitation (ChIP)

Approximately 5 x 10^6^ control and FXS LCLs were harvested and crosslinked with 1% formaldehyde at room temperature for 15 min. Crosslinking was quenched by adding 150 mM glycine and incubating at room temperature for 10 min. Crosslinked cell pellets were lysed in lysis buffer (50 mM Tris-HCl pH 8.0, 10 mM EDTA, 1% SDS) supplemented with protease and phosphatase inhibitor cocktails, and the lysate was diluted with dilution buffer (20 mM Tris-HCl pH 8.0, 150 mM NaCl, 2 mM EDTA, 1% Triton X-100). Chromatin was digested with 200 U MNase (NEB, Cat # M0247S) in the presence of 3 mM CaCl₂ at 37°C for 15 min and sonicated using a Bioruptor (Diagenode) with cycles of 30 s on and 30 s off for 10 cycles. 3.5 µg of fragmented chromatin were subjected to immunoprecipitation with 3 µg of normal mouse IgG (Santa Cruz, Cat # sc-2025) or p-Ser5 Pol II antibody (Abcam, Cat # ab5408) or total Pol II antibody (Santa Cruz, # sc-55492). ChIP DNA was captured using Dynabeads Protein A and G and washed sequentially with Wash buffer A (50 mM Tris-HCl pH 8.0, 140 mM NaCl, 1 mM EDTA pH 8.0, 0.1% SDS, 0.1% sodium deoxycholate), Wash buffer B (50 mM Tris-HCl pH 8.0, 500 mM NaCl, 1 mM EDTA pH 8.0, 1% Triton X-100, 0.1% SDS, 0.1% sodium deoxycholate), Wash buffer C (20 mM Tris-HCl pH 8.0, 250 mM LiCl, 1 mM EDTA pH 8.0, 0.5% NP-40, 0.5% sodium deoxycholate), and twice with TE buffer (10 mM Tris-HCl pH 8.0, 1 mM EDTA). Samples were reverse crosslinked by incubation overnight at 65°C in the presence of RNase A (Invitrogen, Cat # AM2270), followed by protein digestion with Proteinase K at 45°C for 2 h. Chromatin was purified using QIAquick PCR Purification Kit and analyzed by qPCR.

### RNA-seq data processing and analysis

FASTQ files from RNA-seq datasets of FXTAS cerebellum (GSE283760, (23)), FXS white blood cells (GSE202179, (10)), and purified human astrocytes (GSE73721, (24)), were processed using the Via Foundry platform (25). Reads were aligned to the reference genome using STAR (v2.7.9a), and transcript-level quantification was performed with Salmon (v1.9.0). Differential expression analysis was performed using DESeq2 (version 1.40.2). Alternative splicing analysis was conducted with rMATS-turbo (version 4.1.2). mRNA-seq reads and Ribo-seq footprints from LCLs (26) were obtained from GWIPS-viz, and visualized using Gviz (27) in R. GO-term enrichment analysis of the skipped exon events was performed using clusterProfiler.

### Sequence alignment, RNA-binding motif prediction, and CLIP-seq analysis

DNA sequence alignment was performed using MEGA (28). The potential PTBP1 and 2, TDP-43, and SFPQ binding motifs were predicted using RBPmap (29). PTBP1 and TDP-43 binding sites within *FMR1* were obtained from the POSTAR3 database (30), which aggregates ENCODE eCLIP-seq datasets. For PTBP2 eCLIP analysis, FASTQ files from human cortex CLIP-seq data (GSE206661, (31)) were processed and BAM files were generated using ViaFoundry and peaks were visualized using Gviz.

## Results

### Molecular and functional characterization of *FMR1-217*

In typically developing (TD) individuals, the *FMR1* gene contains <55 CGG repeats that allows normal transcription and translation of *FMR1* RNA. In pre-mutation carriers with CGG repeat lengths of 55-200, the *FMR1* locus is transcribed at an elevated level but FMRP is reduced. In FXS, *FMR1* has >200 CGG repeats, leading to gene methylation, transcriptional silencing, and loss of FMRP. Surprisingly in most FXS individuals, *FMR1* transcription persists and gives rise to an aberrantly spliced isoform, *FMR1-217*, consisting of exon 1 fused to a pseudo-exon derived from intron 1 (Fig. 1A).

**Figure 1.**
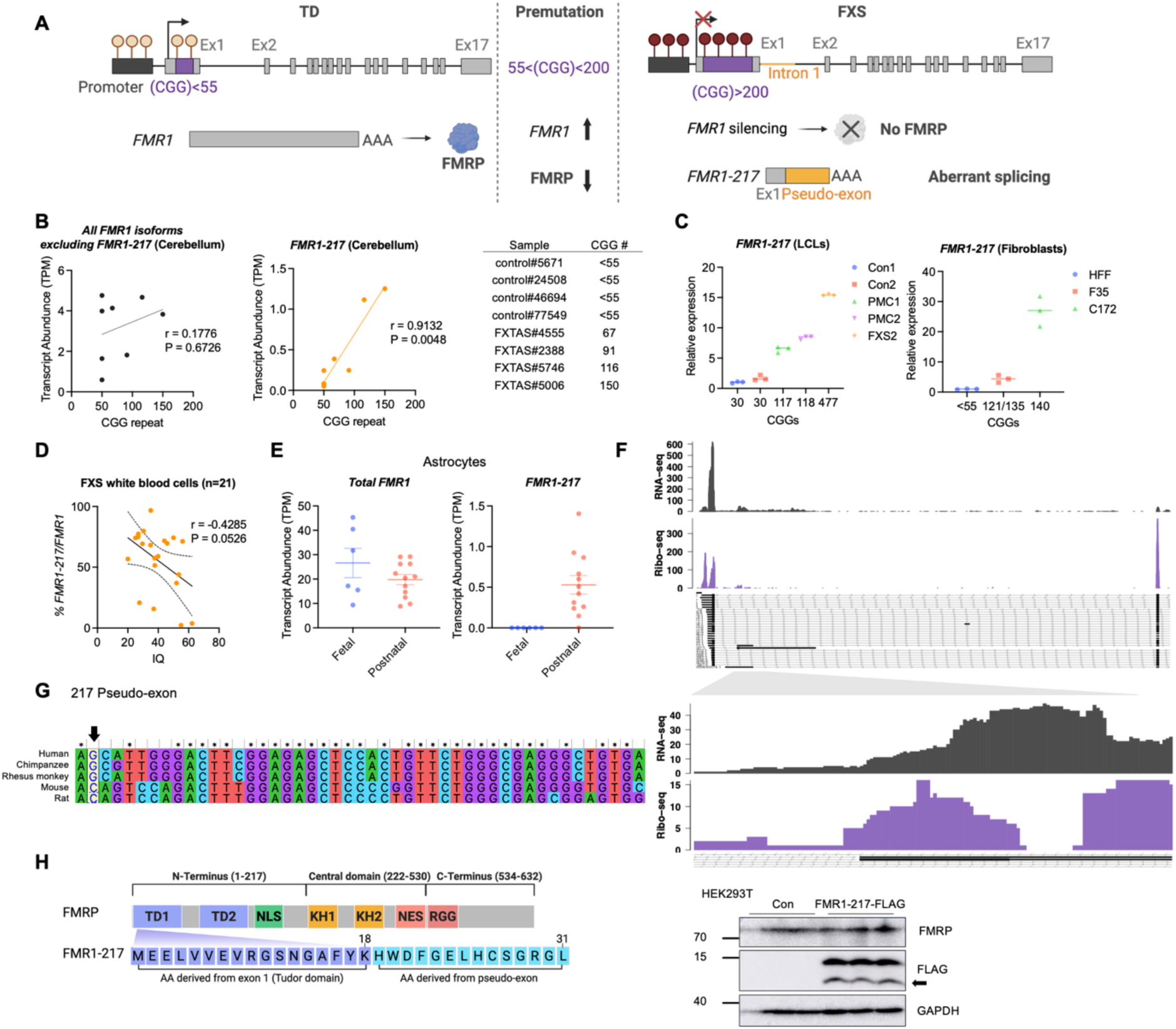
Characterization of *FMR1-217*. (A) Schematic summarizing *FMR1*, *FMR1-217* mRNA and FMRP expression with varying CGG repeat lengths. Typically developing (TD) individuals express *FMR1* mRNA and FMRP protein. Premutation carriers have elevated *FMR1* mRNA but reduced FMRP. FXS individuals lack *FMR1* and FMRP but express the *FMR1-217* transcript. (B) Normalized transcript expression levels (TPM) of *FMR1* (excluding *FMR1-217*) and *FMR1-217* from RNA-seq of human postmortem cerebellum (4 controls, 4 premutation carriers that developed FXTAS). CGG repeat lengths for premutation carriers are listed in the table. The repeat length for control samples is set to 50. RNA-seq data from Yang et al. (23). (C) (Left) *FMR1-217* mRNA levels in human lymphoblastoid cell lines (LCLs) from two controls (Con1 and Con2, 30 CGGs), two premutation carriers (PMC1: 117 CGGs, PMC2: 118 CGGs), and one FXS patient (FXS2, 477 CGGs). (Right) *FMR1-217* mRNA levels in human fibroblast line from control (HFF, <55 CGGs) and two premutation carriers (F35: 35/121/135 CGGs, C172: 140 CGGs). n = 3. (D) Correlation between the percentage of *FMR1-217* relative to total *FMR1* and IQ (n = 21), based on normalized TPM values from FXS white blood cells RNA-seq. Data from Shah et al. (10). The dotted lines in the figure refer to 95% confidence interval. (E) Transcript abundance (TPM) of total *FMR1* and *FMR1-217* in fetal and postnatal human cortical astrocytes. RNA-seq data from Zhang et al. (24). n = 6 for fetal samples and n = 12 for postnatal samples. (F) Ribo-seq and RNA-seq coverage across the *FMR1* exon 1-exon 2 region (left) and the *FMR1-217* pseudo-exon (right). RNA-seq read density (top, gray) and ribosome profiling (Ribo-seq) signal (bottom, purple) are shown. RNA-seq and ribosome profiling data from Battle et al. (26) are obtained from GWIPS-viz (n = 72). Below is a western blot for *FMR1-217* 3x FLAG upon transfection into HEK293T cells. The upper band is likely to be a dimer of the peptide. (G) Sequence conservation across six mammals (human, chimpanzee, rhesus monkey, mouse, rat) is shown below, with the critical splice-site G nucleotide (arrow), absent in mouse and rat. (H) Schematic representation of the domain architecture of FMRP and the predicted *FMR1-217* peptide. TD (Tudor/Agenet domain), NLS (nuclear localization signal), KH (RNA-binding domain), NES (nuclear export signal), and RGG (RNA-binding domain) are indicated. The *FMR1-217* peptide consists of 18 amino acids derived from exon 1, corresponding to part of the Tudor domain, and 13 amino acids encoded by the pseudo-exon.

Our previous study (10) reported that *FMR1-217* is not detected in skin biopsy-derived fibroblasts from a premutation carrier with 98 CGG repeats but is present in an individual with 140 CGG repeats suggesting a CGG repeat-length dependent threshold for *FMR1-217* expression. To more precisely define this threshold required for *FMR1-217* expression, we reanalyzed published RNA-seq data from human cerebellum samples spanning a range of CGG repeat length (67, 91, 116, and 150 repeats) obtained from normal and FXTAS patients (23). *FMR1-217* expression was low at 91 or fewer CGGs but similarly high at 116 or 150 CGGs, indicating a CGG length dependency to produce *FMR1-217. FMR1-217* expression was strongly correlated with CGG repeat expansion (r = 0.9132, p = 0.0048) (Fig. 1B). In contrast, *FMR1* isoforms excluding *FMR1-217* showed a modest positive trend with CGG repeat length, indicating that this strong CGG length dependency is specific to *FMR1-217*. We quantified the *FMR1-217* RNA level in lymphoblastoid cells (LCLs) derived from typically developing (TD) individuals, pre-mutation carriers, and a FXS individual (Fig. 1C, left). Consistent with RNA-seq data from FXTAS brain samples (Fig. 1B) and previous findings (10), *FMR1-217* expression was detected in both premutation carriers with 117 and 118 repeats and FXS but not two TD individuals with 30 CGGs. To further validate this relationship, we quantified *FMR1-217* levels in two FXTAS patient-derived fibroblast lines relative to control fibroblasts: one with a mosaic 32, 121, and 135 repeats, and another with 140 CGG repeats. *FMR1-217* levels showed an increasing trend in the mosaic line and were strongly increased in the fibroblast line carrying 140 CGG repeats compared to control, consistent with a repeat length dependent increase in *FMR1-217* expression (Fig. 1C, right). These data confirm a positive relationship between CGG repeat size and *FMR1-217* mis-splicing.

Both total *FMR1* and *FMR1-217* levels have been reported to correlate negatively with IQ in FXS individuals (10). However, because absence of total *FMR1* itself influences IQ, it is unclear whether *FMR1-217* makes an independent contribution to cognitive outcome. To isolate the potential effect of *FMR1-217* from that of total *FMR1*, we reanalyzed a published RNA-seq dataset from white blood cells of FXS individuals, restricting the analysis to samples with detectable *FMR1* expression (n = 21) to avoid confounding by complete transcriptional silencing. We calculated the percentage of *FMR1-217* relative to total *FMR1* transcript as a measure of relative mis-splicing and examined its association with IQ (Fig. S1A). The *FMR1-217* percentage was negatively correlated with IQ (r = −0.4285, p = 0.0526) (Fig. 1D), while total *FMR1* showed no correlation (Fig. S1B). This suggests that a higher proportion of the mis-spliced isoform is associated with poorer cognitive outcomes in FXS independently of total *FMR1* abundance.

To determine whether *FMR1-217* expression is developmentally regulated, we reanalyzed RNA-seq data from Zhang et al. (24), who developed an immunopanning-based purification method to isolate human neurons, glial cells, and vascular cells. We compared *FMR1-217* expression in human astrocytes at two developmental stages: fetal and postnatal. Although total *FMR1* expression levels were similar between stages, fetal cortical astrocytes showed no detectable *FMR1-217* expression (Fig. 1E), whereas postnatal astrocytes showed varying but elevated levels of *FMR1-217*, suggesting developmental regulation of this isoform.

To further characterize *FMR1-217*, we analyzed GWIPS-viz ribosome profiling and RNA-seq data (26,32) from human LCLs samples. Ribosome footprints were detected within the pseudo-exon region, indicating that *FMR1-217* is likely translated into a short 31 amino acid peptide (Fig. 1F). Indeed, a reporter RNA encoding the *FMR1-217* peptide followed by multiple FLAG epitopes is translated upon cell transfection, without affecting endogenous FMRP protein levels. Two bands were observed, which may be *FMR1-217* dimers as FMRP is known to dimerize via its amino terminal region (33) (Fig. 1F). However, we were unable to detect the predicted 31 amino acid peptide by mass spectrometry, consistent with reports that only a small fraction of peptides predicted from ribosome footprints on non-canonical regions (uORFs, introns, 3’UTRs) are validated by mass spectrometry (34). Sequence conservation analysis revealed that the splice sites associated with *FMR1-217* are highly conserved across three primate species, but not in mice or rats, which both lack a critical guanosine at the 3’ splice site (Fig. 1G, arrow). This observation as well as the absence of CGG expansion in mice, explains why *FMR1-217* isoform is not expressed in this animal model. The *FMR1-217* peptide contains 18 amino acids derived from exon 1, which correspond to a portion of the Tudor domain that mediates protein–protein interactions and chromatin binding, followed by 13 additional amino acids encoded by the pseudo-exon (Fig. 1H).

### CGG repeat expansions promote R-loop formation and *FMR1-217* mis-splicing

To assess the functional consequences of *FMR1-217* expression, we selected six unmethylated full mutation (UFM) FXS samples with comparable total *FMR1* expression levels from a published white blood cells RNA-seq dataset. Samples were divided into a *FMR1-217*-high group (>70% of total *FMR1*) and a *FMR1-217*-low group (<21%) (Fig. 2A). Principal component analysis revealed clear separation between the two groups (Fig. S2A), and differential expression analysis demonstrated that the proportion of *FMR1-217* is associated with distinct gene expression changes (Fig. 2B). Notably, *PIF1*, a DNA helicase known to resolve DNA-RNA hybrids, was significantly upregulated in the *FMR1-217*-high group, suggesting an increased R-loop burden in these cells. Consistent with this, *PIF1* mRNA is elevated in one premutation carrier and in the FXS LCLs (Fig. 2C). However, because a reduction of PIF1, a DNA helicase, is linked to R-loop formation (35,36), we examined the levels of this protein in premutation carrier (PMC) and FXS2 cells. PIF1 was markedly reduced in FXS2 cells despite elevated *PIF1* mRNA (Fig. 2D), suggesting that reduced PIF1 protein may indeed contribute to R-loop accumulation in FXS. There is often a disconnect between RNA levels and their encoded proteins, a phenomenon that has been shown frequently in FXS model mice using ribosome profiling or SILAC (Stable Isotope Labeling by Amino Acids in Cell Culture) labeling of protein (e.g.,(37–39)). We also observed widespread alternative splicing changes independent of total *FMR1* levels (Fig. S2B, S2C), including enrichment for cillum assembly, autophagy, glycerophospholipid metabolism, and histone modification, as well as alternative splicing regulators themselves (e.g., PTBP1, HNRNPs, SRSF4), suggesting these regulators may contribute to even broader splicing changes. Together, these findings implicate R-loop-associated mis-splicing as a potential mechanism linking CGG repeat expansion to *FMR1-217* expression.

**Figure 2.**
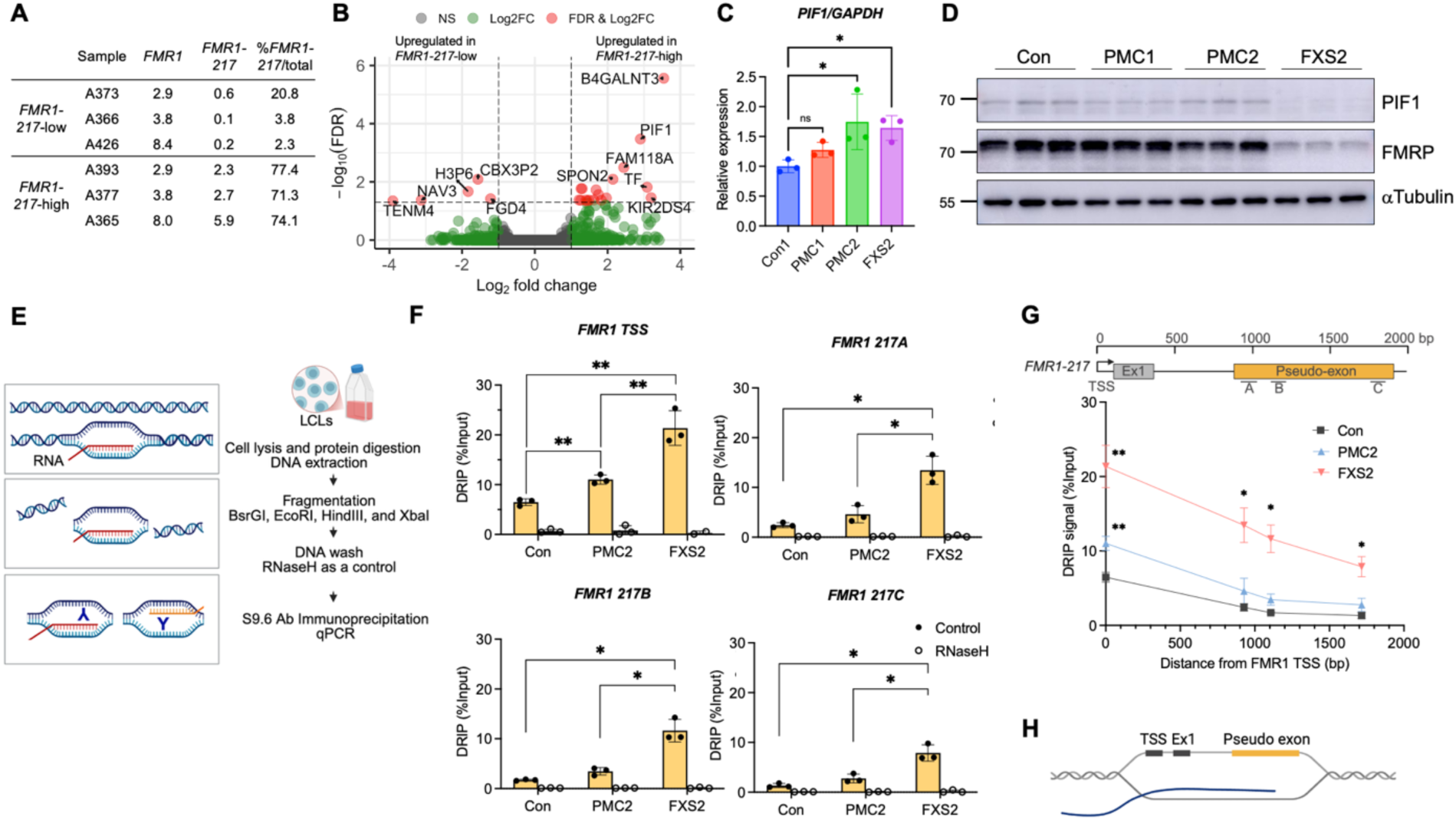
R-loop accumulation from *FMR1* TSS to the *FMR1-217* pseudo-exon. (A) Table summarizing TPMs of *FMR1* and *FMR1-217*, and the proportion of *FMR1-217* relative to total *FMR1* in *FMR1-217*-low and -high groups. (B) Volcano plot of differential gene expression between *FMR1-217*-low and -high groups. Significantly altered genes (FDR < 0.05 and |log₂ fold change| > 1) are highlighted in red. (C) *PIF1* mRNA levels in LCLs (n = 3). Error bars represent SD. Statistical significance was determined by one-way ANOVA. *P < 0.05 (D) Immunoblot of PIF, FMRP, and α-Tubulin. (E) Schematic of DRIP-qPCR workflow. (F) DRIP-qPCR analysis of R-loop signal at the *FMR1* TSS and within the pseudo-exon (*FMR1-21A-C*) in control (Con), premutation carrier (PMC2), and FXS (FXS2) LCLs. Samples were analyzed with (open circles) or without RNaseH (filled circles). Error bars represent SD, n = 3. Statistical significance was determined by two-tailed Student’s t-test. *P < 0.05, **P < 0.01, ***P < 0.001. (G) Schematic showing the primer locations across the *FMR1-217* region and corresponding DRIP signal plotted relative to distance from the *FMR1* TSS. Error bars represent SD, n = 3. Statistical significance was determined by two-tailed Student’s t-test. *P < 0.05, **P < 0.01, ***P < 0.001. (H) Model illustrating R-loop formation at the *FMR1* locus.

R-loops are enriched at CGG repeat expansions within the *FMR1* locus (19). Consistent with this, elevated *PIF1* RNA but reduced PIF1 protein expression in the *FMR1-217-high* group may increase DNA-RNA hybrid formation. We therefore hypothesized that R-loop accumulation near expanded CGG repeats at the *FMR1* locus may contribute to *FMR1-217* mis-splicing.

To map R-loop formation across *FMR1*, we performed DNA-RNA immunoprecipitation (DRIP)-qPCR. DNA was fragmented with restriction enzyme cocktails and R-loop-containing regions were immunoprecipitated using the S9.6 antibody. To confirm the specificity of immunoprecipitation signal, chromatin was treated with RNase H, which specifically degrades RNA in DNA-RNA hybrids (Fig. 2E). R-loops were enriched at the transcription start site (TSS), consistent with a previous report (19), and were detected in both premutation carrier (PMC2) and FXS (FXS2) cells relative to control (Fig. 2F). R-loop distribution across *FMR1-217* was visualized by plotting DRIP signal intensity relative to the distance from the TSS. R-loops were more extensive and larger in FXS2 than in PMC2 (Fig. 2G), suggesting that CGG repeat expansion enhances R-loop formation in a repeat length-dependent manner, spanning from the TSS into the pseudo-exon (Fig. 2H). *RPL13A* and *EGR1* were used as positive and negative controls, respectively (Fig. S2D).

### ASO treatment reduces *FMR1-217* without affecting R-loop formation

We showed that antisense oligonucleotides (ASOs 713,714) targeting the pseudo-exon reduced *FMR1-217* mRNA expression and restored FMRP (10). To determine whether ASO-mediated suppression of mis-splicing involves R-loop resolution, we treated control and fully methylated FXS (FXS1) LCLs with ASOs, prior to demethylation using nucleoside analog 5-AzadC for 5 days to reactivate *FMR1* transcription, as previously described (10) (Fig. 3A). Because 5-AzadC reactivates transcription of the entire *FMR1* locus, this treatment increased levels of both full-length *FMR1* and *FMR1-217* RNA (Fig. 3A). Western blot analysis confirmed that 5-AzadC reactivated FMRP and that ASO treatment further increased its expression (Fig. 3B). ASO treatment of cells whose transcription was stimulated by 5-AzadC reduced *FMR1-217* RNA levels, although not significantly (Fig. 3C). The ASOs reduced *FMR1-217* levels, but DRIP-qPCR revealed no change in R-loop signal, indicating that the ASOs reduced *FMR1-217* through an R-loop-independent mechanism (Fig. 3D-E). R-loops in *RPL13A* and *EGR1* were detected as positive and negative controls, respectively (Fig. 3F).

**Figure 3.**
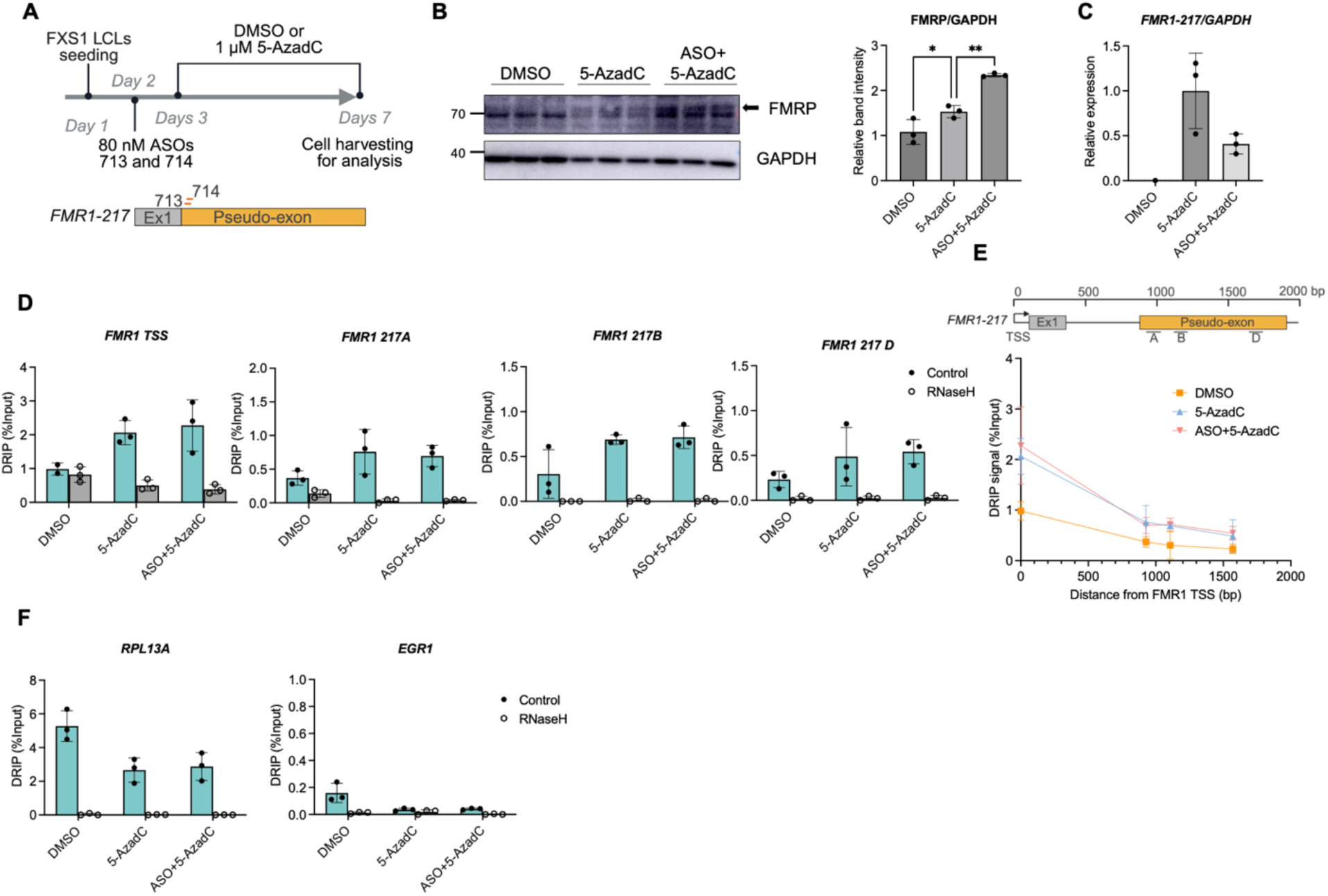
Effect of ASO treatment on *FMR1-217* and R-loop formation in FXS LCLs. (A) Experimental schematic showing ASO and 5-AzadC treatment in fully silenced FXS (FXS1) LCLs. (B) Immunoblot of FMRP and GAPDH. The FMRP band is indicated by an arrow. Quantification of FMRP levels normalized to GAPDH is shown on the right. Error bars represent SD (n = 3). Statistical significance was determined by one-way ANOVA. *P < 0.05, **P < 0.01. (C) *FMR1-217* mRNA levels following 5-AzadC and/or ASO treatment. Error bars represent SD (n = 3). (D) DRIP-qPCR analysis showing changes in R-loop signal. Samples were analyzed with (open circles) or without RNaseH (filled circles). Error bars represent SD, n = 3. (E) Schematic of primer locations used for R-loop detection with DRIP signal from the TSS through the pseudo-exon. Corresponding DRIP signals were plotted relative to distance from the *FMR1* TSS. Error bars represent SD, n = 3. (F) DRIP-qPCR analysis of R-loop enrichment at *RPL13A* and *EGR1* in FXS1 LCLs treated with DMSO or 5-AzadC and/or ASOs with or without RNase H treatment. Error bars represent SD (n = 3).

### Pol II elongation is impaired at the 5’ region of *FMR1* in FXS cells

Co-transcriptional splicing is sensitive to RNA polymerase II (Pol II) elongation kinetics where local pausing or stalling can alter splice site recognition (40). To investigate whether altered Pol II dynamics at the 5’ region of *FMR1* contribute to *FMR1-217* pseudo-exon inclusion in FXS cells, we performed a DRB washout assay. DRB (5,6-dichloro-1-β-D-ribofuranosylbenzimidazole) inhibits CDK9 within the Positive Transcription Elongation Factor (P-TEFb) complex, preventing phosphorylation of the Pol II C-terminal domain (CTD) at serine 2, and thereby blocking productive elongation. Control and FXS LCLs were treated with DRB for 3 hours, followed by washout and recovery in fresh media for 5, 10, and 15 minutes (Fig. 4A). *FMR1* transcripts were quantified by qPCR using primer pairs targeting the exon 1-intron 1 junction, the intron 1-pseudo-exon junction, and the intron 2-exon 2 junction (Fig. 4B). Following DRB removal, *FMR1* transcript levels increased over time in both control and FXS cells, indicating recovery of transcription.

**Figure 4.**
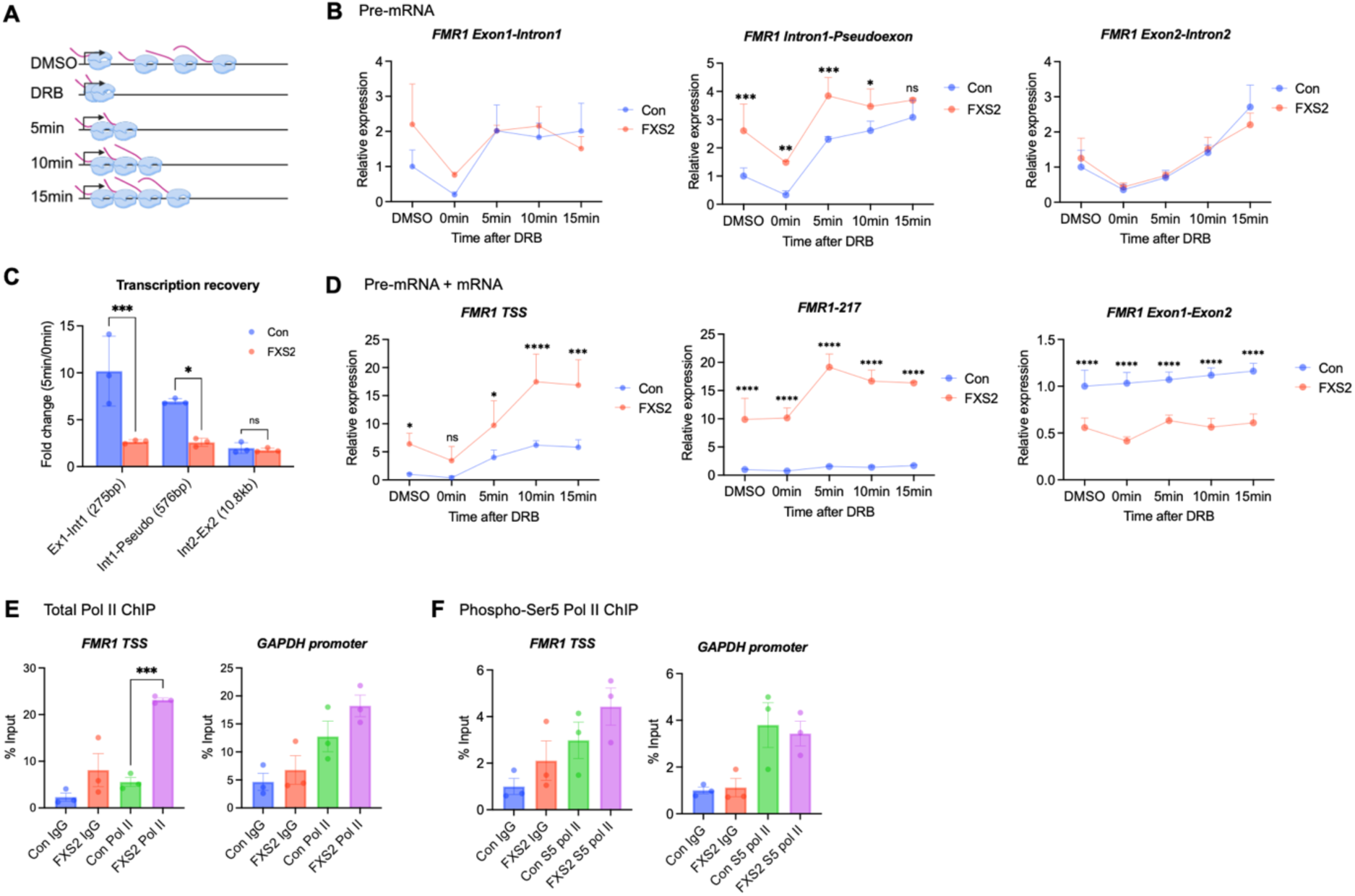
Transcription perturbation at the 5’ region of the *FMR1*. (A) Schematic of DRB washout assay. Control and FXS2 LCLs were treated with DMSO or DRB for 3 h and collected 5, 10, and 15 min after DRB removal. (B) Comparison of *FMR1* pre-mRNA levels measured in control and FXS2 LCLs treated with or without DRB, using primers targeting the exon 1-intron 1, intron 1-pseudo-exon junction, and exon 2-intron 2 junction. Error bars represent SD (n = 3). Statistical analysis was performed using two-way ANOVA followed by Turkey’s multiple comparisons test. *P <0.05, **P < 0.01, ***P < 0.001, ****P < 0.0001. (C) Transcription recovery following DRB washout. The fold change in signal for each amplicon was calculated as the 5 min/ 0 min ratio. Error bars represent SD (n = 3). (D) Relative mRNA expression levels of *FMR1* at the TSS, mRNA expression levels of *FMR1-217* and full-length *FMR1* in control and FXS2 LCLs treated with or without DRB. All primer sets detect both pre-mRNA and mature mRNA. Error bars represent SD (n = 3). Statistical analysis was performed using two-way ANOVA followed by Turkey’s multiple comparisons test. *P <0.05, **P < 0.01, ***P < 0.001, ****P < 0.0001. (E) ChIP-qPCR assay of total Pol II. Error bars represent SD (n = 3). Statistical significance was determined by two-tailed Student’s t-test, ***P < 0.001. (F) ChIP-qPCR assay of phospho-Ser5 Pol II. Chromatin was immunoprecipitated with mouse IgG or phospho-Ser5 RNA Pol II antibody and analyzed by qPCR using primer pairs targeting the *FMR1* TSS and GAPDH promoter region. Error bars represent SD (n = 3).

Because FXS cells showed markedly higher baseline levels than control cells at 0 min, the recovery slopes are not directly comparable. We therefore quantified recovery as fold change (5 min/0 min) for each amplicon (Fig. 4C). FXS cells exhibited lower transcription recovery at the exon 1-intron 1 and the intron 1-pseudo-exon junction, but not at the intron 2-exon 2 junction, suggesting that transcription perturbation is localized to the 5’ region of the *FMR1* containing the pseudo-exon.

In FXS cells, *FMR1-217* RNA levels were elevated at the TSS and intron 1-pseudo-exon junction at the 0-minute timepoint and increased further within 5 minutes following washout (Fig. 4D). The elevated mRNA at the TSS could reflect enhanced transcription or accumulation of promoter-proximal pausing (41,42). To directly test whether Pol II accumulates at the *FMR1* TSS in FXS, we performed chromatin immunoprecipitation (ChIP)-qPCR for total Pol II. Control and FXS LCLs were crosslinked and digested with micrococcal nuclease (MNase) to generate chromatin fragments (Fig. S3A), followed by immunoprecipitation with either IgG control or total Pol II antibody. We found significantly increased total Pol II occupancy at the *FMR1* TSS in FXS cells relative to control, with no change at the *GAPDH* promoter (Fig. 4E). To determine whether this reflects an increase in engaged, promoter-proximal paused polymerase, we performed ChIP for phospho-Ser5 Pol II and found no significant difference in occupancy between control and FXS cells (Fig. 4F). Together, these results indicate that Pol II accumulates at the *FMR1* TSS in FXS cells in a hypophosphorylated, non-actively initiating state.

### CPT induced Pol II stalling promotes *FMR1-217* mis-splicing

To directly test whether experimentally induced Pol II stalling influences *FMR1* splicing, we treated cells with camptothecin (CPT), a topoisomerase I inhibitor that stabilizes the Top1-DNA cleavage complex (Top1cc) and creates a physical barrier to Pol II progression along the gene body (Fig. 5A). Although both DRB and CPT affect Pol II transcription, they act through distinct mechanisms and have differential effects on alternative splicing (43). CPT treatment has also been also reported to induce accumulation of Ser5-phosphorylated Pol II as a consequence of transcriptional arrest (44).

**Figure 5.**
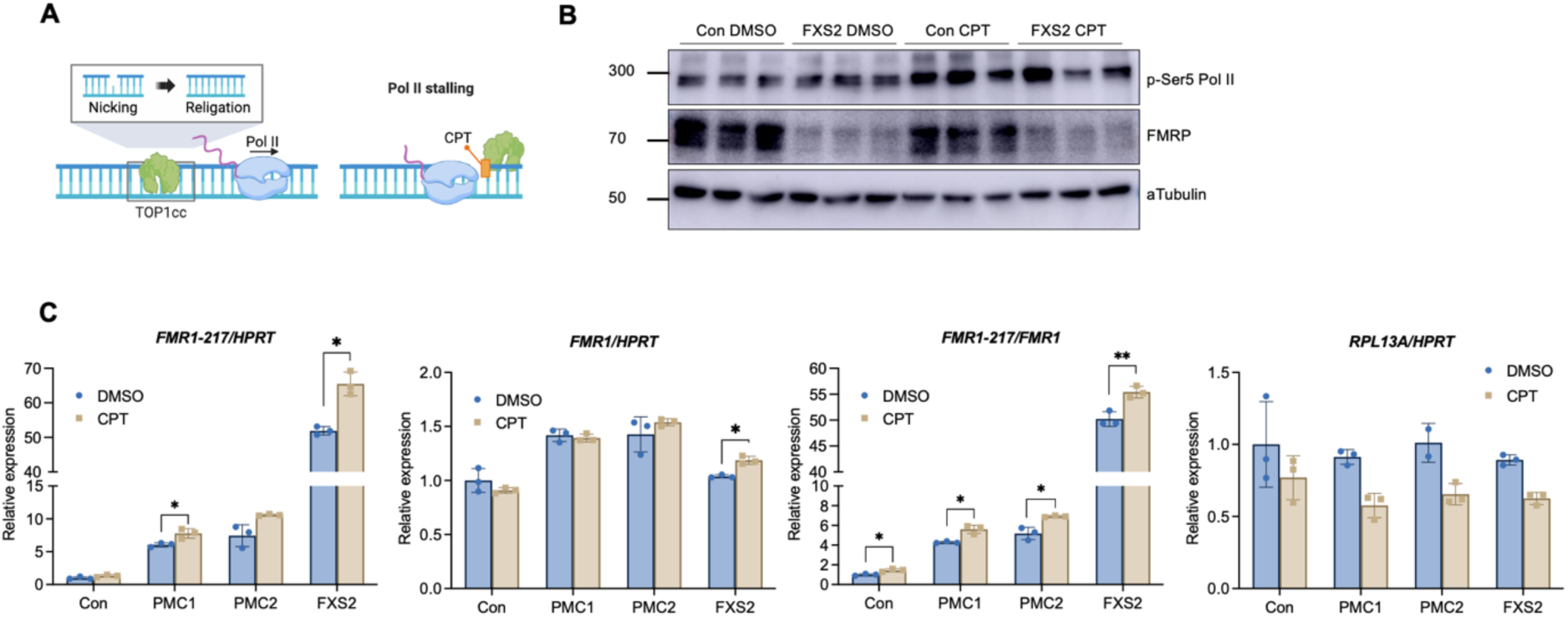
Pol II stalling increases *FMR1-217* mis-splicing. (A) Schematic illustrating the mechanism by which CPT induces Pol II stalling. (B) Immunoblot analysis of phospho-Ser5 RNA polymerase II (p-Ser5 Pol II), FMRP, and α-Tubulin in control and FXS2 LCLs treated with DMSO or CPT. α-Tubulin was used as a loading control. (C) *FMR1-217* and full-length *FMR1* mRNA levels in control, PMC1, PMC2, and FXS2 LCLs following 1 h treatment with DMSO or CPT. *FMR1* and *FMR1-217* levels were normalized to *HPRT*, and *FMR1-217* levels are also shown relative to full-length *FMR1*. *RPL13A* mRNA was measured as a positive control. Error bars represent SD (n = 3). Statistical significance was determined by two-tailed Student’s t test. *P < 0.05, **P < 0.01, ***P < 0.001.

Control, premutation carrier, and FXS LCLs were treated with CPT for 1 hour. Increased Ser5 phosphorylation of the Pol II CTD was confirmed by Western blotting (Fig. 5B), validating transcription perturbation. Consistent with global transcription inhibition, expression of the control gene *RPL13A* decreased by approximately 30% following CPT treatment (Fig. 5C). In contrast, full-length *FMR1* levels showed only a modest change, whereas *FMR1-217* levels exhibited a strong increase in FXS2 cells. Normalization of *FMR1-217* to full-length *FMR1* transcript revealed a significant increase in the relative abundance of the mis-spliced isoform following CPT treatment (Fig. 5C), indicating that Pol II stalling selectively promotes *FMR1-217* mis-splicing relative to canonical splicing. Together with the DRB washout findings, these results suggest that perturbation of co-transcriptional splicing contributes to *FMR1-217* generation.

### PTBP1 regulates *FMR1-217* mis-splicing in FXS fibroblasts

To identify splicing factors that may contribute to *FMR1-217*, we analyzed the pseudo-exon region using RBPmap (29) and identified predicted binding sites for polypyrimidine tract-binding proteins (PTBPs) (Fig. 6A). PTBP1 is a well-characterized splicing repressor that binds CU-rich sequences and suppresses inclusion of cryptic exons (45). To determine whether PTBP1 directly interreacts with the *FMR1* pre-mRNA, we examined published PTBP1 eCLIP-seq data from HepG2 and K562 cells using POSTAR3 database (30). We identified PTBP1 binding sites within *FMR1* intron 1 downstream of the pseudo-exon, and within intron 3. Because PTBP2 shares a highly similar binding motif with PTBP1 and is the predominant paralog expressed in brain, we also reanalyzed published human cortex PTBP2 CLIP-seq data (31). CLIP peaks were detected within the *FMR1* CGG repeat region as well as at three additional sites in intron 1 downstream of the *FMR1-217* pseudo-exon (Fig. 6B), several of which overlap with the PTBP1 intron 1 sites. Together, these data indicate that PTBP1 and PTBP2 may both bind directly to *FMR1*.

**Figure 6.**
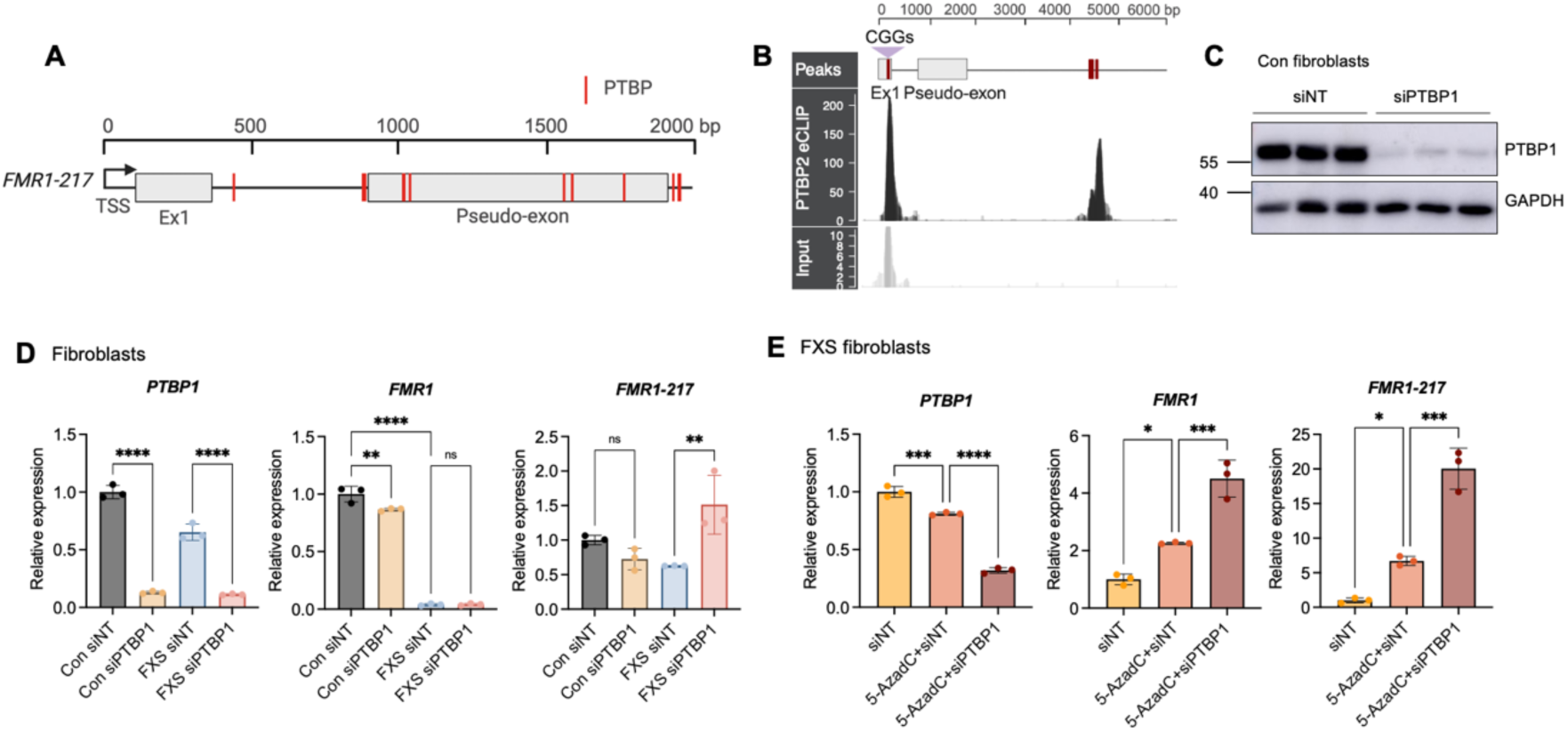
PTBP1 regulates *FMR1-217* expression in FXS fibroblasts. (A) Schematic of predicted PTBP binding motifs within *FMR1-217*. (B) PTBP2 enhanced CLIP (eCLIP) read coverage in human cortex, with size matched input read coverage (n =3, overlaid). Peaks located within the CGG repeat and intron 1 are highlighted in red. The *FMR1* gene model (exon 1 and pseudo-exon) is shown for reference. (C) Immunoblot showing PTBP1 knockdown in control (Con) fibroblasts. GAPDH was used as a loading control. (D) mRNA levels of *PTBP1*, *FMR1*, and *FMR1-217* in Con and FXS fibroblasts treated with non-targeting siRNA (siNT) or siPTBP1. Error bars represent SD (n =3). Statistical significance was determined by one-way ANOVA. **P < 0.01, ****P < 0.0001. (E) FXS fibroblasts were treated with 5-AzadC for 7 days to reactivate *FMR1* expression. siRNAs were transfected on day 4. mRNA levels of *PTBP1*, *FMR1*, and *FMR1-217* are shown. Error bars represent SD (n =3). Statistical significance was determined by one-way ANOVA. *P < 0.05, ***P < 0.001, ****P < 0.0001.

To investigate whether PTBP1 regulates *FMR1-217* splicing, experiments were performed in fibroblasts rather than LCLs, as siRNA-mediated knockdown is technically challenging in suspension cultures due to low transfection efficiency. siRNA targeting PTBP1 was transfected into fibroblasts derived from TD individuals (Con) and 81%-methylated mosaic FXS patients (FXS). Western blot analysis confirmed efficient PTBP1 knockdown (Fig. 6C), and qPCR showed a corresponding reduction in *PTBP1* mRNA levels in both Con and FXS fibroblasts. PTBP1 depletion significantly increased *FMR1-217* expression (Fig. 6D).

Unlike FXS2 LCLs, FXS fibroblasts do not express either *FMR1* or *FMR1-217,* Consequently, FXS cells were treated with 5-AzadC for five days to reactivate *FMR1* transcription. Demethylation induced expression of both full-length *FMR1* and *FMR1-217*. Subsequent PTBP1 knockdown further increased full-length *FMR1* expression approximately 2-fold and *FMR1-217* expression approximately 3-fold, suggesting that PTBP1 suppresses *FMR1-217* pseudo-exon inclusion (Fig. 6E).

### TDP-43 regulates both *FMR1* and *FMR1-217* production in FXS fibroblasts

There are several TDP-43 predicted binding sites in *FMR1-217* (Fig. S4A) identified by RBPmap, although there are no TDP-43 binding sites within *FMR1* were found in the POSTAR3 database (30). TDP-43 is an RNA-binding protein involved in splicing regulation and RNA processing and its dysfunction is linked to aberrant cryptic exon inclusion in neurodegenerative disease. In motor neurons, TDP-43 represses cryptic splicing of *STMN2*, a gene required for axonal regeneration following injury. In amyotrophic lateral sclerosis (ALS), nuclear depletion of TDP-43 leads to activation of a cryptic splice site within intron 1 of *STMN2,* resulting in truncated transcripts and loss of functional protein (22).

To investigate the role of TDP-43 in *FMR1* splicing, we depleted it in control and FXS fibroblasts using siRNA. Under basal conditions, TDP-43 depletion did not significantly affect expression of either full-length *FMR1* or *FMR1-217*, while its known target, truncated *STMN2*, was upregulated, confirming that TDP-43 depletion was functional (Fig. S4B-D). Following *FMR1* reactivation with 5-AzadC, TDP-43 depletion significantly reduced expression of both full-length *FMR1* and *FMR1-217* (by 67% and 76%, respectively), suggesting that TDP-43 broadly affects *FMR1* rather than selectively promoting *FMR1-217* splicing (Fig. S4E).

### PTBP1/PTBP2 regulate *FMR1-217* splicing in a differentiation stage-dependent manner

We examined the roles of PTBP1 in splicing regulation in both TD and unmethylated full-mutation FXS iPSCs to assess a cell type dependent splicing regulation while excluding potential off-target effects caused by 5-AzadC treatment.

Transfection with siRNA targeting PTBP1 efficiently reduced both PTBP1 protein and mRNA levels (Fig. 7A-B). Unlike in fibroblasts, PTBP1 knockdown in FXS iPSCs did not alter full-length *FMR1* mRNA expression but significantly reduced *FMR1-217* levels (Fig. 7B), indicating that PTBP1 promotes pseudo-exon inclusion. Although PTBP1 is broadly recognized as a splicing repressor, its function as a splicing activator has been reported to depend on its binding position relative to the regulated exon (46).

**Figure 7.**
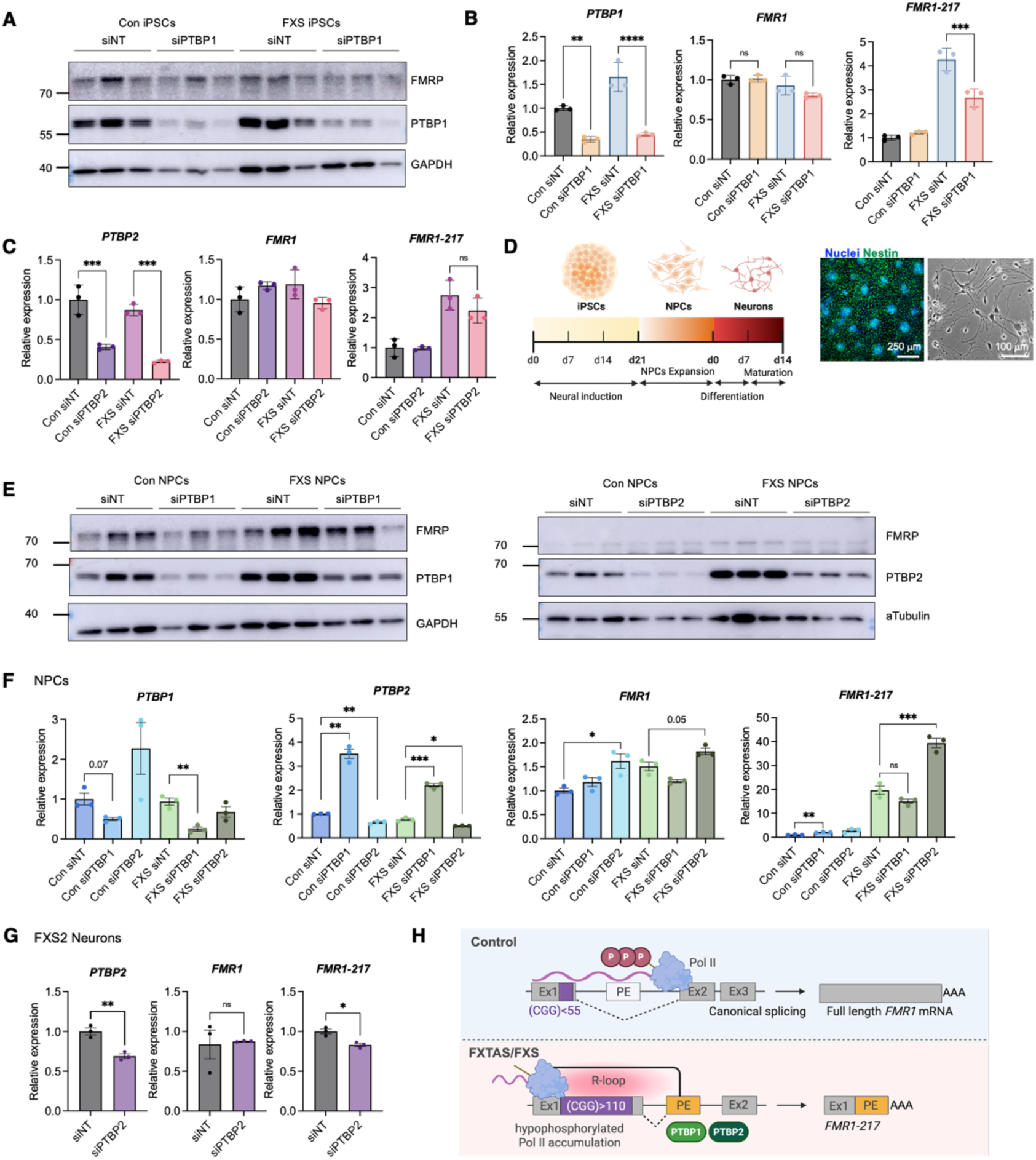
PTBP1 and PTBP2 regulate *FMR1-217* splicing in FXS in a differentiation stage-dependent manner. (A) Immunoblot analysis of FMRP and PTBP1 in control and FXS iPSCs following transfection with siNT or siPTBP1. GAPDH was used as a loading control. (B) Relative mRNA levels of *PTBP1*, *FMR1*, and *FMR1-217* in control and FXS iPSCs treated with siNT or siPTBP1. Error bars represent SD (n =3). Statistical significance was determined by one-way ANOVA (**P < 0.01, ***P < 0.001, ****P < 0.0001). (C) Knockdown of PTBP2 in control and FXS iPSCs did not alter *FMR1-217* levels, although *PTBP2* mRNA was efficiently reduced. Error bars represent SD (n =3). Statistical significance was determined by two-tailed Student’s t-test. ***P < 0.001. (D) Timeline of neural differentiation from iPSCs. Immunofluorescence staining of NPCs for the NPC marker Nestin. Nuclei were stained with Hoechst33342. Scale bar = 200 μM. Bright field image of neurons. Scale bar = 100 μM. (E) Immunoblot analysis of FMRP, PTBP1, and PBP2 in control and FXS NPCs following transfection with siNT or siPTBP1/2. GAPDH or α-Tubulin was used as a loading control. (F) Relative mRNA levels of *PTBP1, PTBP2*, *FMR1*, and *FMR1-217* in control and FXS NPCs treated with siNT or siPTBP1/2. Error bars represent SD (n = 3). Statistical significance was determined by two-tailed Student’s t-test. *P < 0.05, **P < 0.01, ***P < 0.001. (G) Relative mRNA levels of PTBP2, FMR1, and FMR1-217 in FXS neurons transfected with siNT or siPTBP2 on day 5 of maturation. Error bars represent SD (n =3). Statistical significance was determined by two-tailed Student’s t test. *P < 0.05, **P < 0.01. (H) Model of *FMR1-217* mis-splicing. CGG repeat expansion is associated with R-loop formation that may impair Pol II elongation, causing the polymerase to accumulate in a hypophosphorylated state near the 5’ region of *FMR1*. This delays the transition to productive elongation and favors recognition of the downstream, non-canonical pseudo-exon splice site, resulting in *FMR1-217* mis-splicing. Both PTBP1 and PTBP2 are involved in *FMR1-217* mis-splicing in a differentiation stage-dependent manner.

PTBP2, a paralog of PTBP1 with partially redundant splicing functions, is expressed predominantly in neuronal cell types and largely replaces PTBP1 during neuronal differentiation (47). PTBP2 knockdown in FXS iPSCs resulted in a mild, non-significant reduction of *FMR1-217* (Fig. 7C) likely reflecting its low expression in iPSCs. We tested SFPQ, given the presence of a predicted SFPQ binding motif near the pseudo-exon and its role in generating cryptic last exons within long introns and producing cleaved polyadenylated isoforms (48). However, SFPQ knockdown did not affect *FMR1-217* levels (Fig. S5A).

To determine whether PTBP1/PTBP2’s regulation of *FMR1-217* extends to CNS-relevant cell types, we differentiated control and FXS iPSCs into neural progenitor cells (NPCs) and further differentiated FXS NPCs into neurons. NPC identity was confirmed by immunostaining for the NPC marker Nestin, and neuronal differentiation was confirmed by characteristic neuronal morphology (Fig. 7D). In FXS NPCs, PTBP2 knockdown increased *FMR1-217* levels approximately 2-fold with no significant change in full-length *FMR1* (Fig. 7E-F). PTBP1 knockdown significantly increased *PTBP2* mRNA levels, consistent with a known compensatory effect between the two paralogs (45). However, this compensatory increase of *PTBP1* mRNA did not significantly alter *FMR1-217* splicing in FXS NPCs. This suggests that PTBP2 rather than PTBP1 regulates *FMR1-217* splicing in NPCs, where it acts to repress pseudo-exon inclusion.

To determine whether this repressive role of PTBP2 is retained in neurons, we further differentiated FXS NPCs into neurons and performed PTBP2 knockdown during early maturation. In contrast to its repressive effect in NPCs, PTBP2 knockdown in neurons significantly reduced *FMR1-217* splicing (Fig. 7G-H), indicating that PTBP2 switches from a repressor to an activator of pseudo-exon inclusion as NPCs mature into neurons.

We further examined the role of TDP-43 across iPSCs, NPCs, and neurons. TDP-43 knockdown reduced *FMR1-217* in both iPSCs and NPCs. Full-length *FMR1* showed a decreasing trend in iPSCs and was significantly reduced in NPCs, suggesting a general effect of TDP-43 on *FMR1* transcription rather than specific regulation of pseudo-exon splicing (Fig. S5B-E). In immature neurons, TDP-43 knockdown did not significantly alter either *FMR1* or *FMR-217* mRNA levels (Fig. S5F).

Collectively, these findings indicate that PTBP1 and PTBP2 regulate *FMR1-217* splicing in a cell type- and differentiation stage-dependent manner, switching between activating and repressive roles from iPSCs through neurons.

## Discussion

Human *FMR1* is alternatively spliced into as many as 49 isoforms, 30 of which are detected almost exclusively in pre-mutation carriers suggesting they may be related to FXTAS pathology (49). Although *FMR1* is ostensibly silent in FXS, we detected 15 isoforms at varying levels in WBCs from individuals with this disorder. One isoform, *FMR1-217,* composed of exon 1 spliced to a pseudo-exon in intron 1, could be rescued by splice-switching ASOs that in turn restored FMRP to normal levels. These observations, the negative correlation between *FMR1-217* and IQ (10), and a possible treatment of FXS with an ASO therapeutic caused us to focus on this isoform and how it is generated.

We further demonstrated that *FMR1-217* was negatively correlated with IQ independent of total *FMR1* levels, suggesting that *FMR1-217* mRNA, or the peptide it encodes, may have a deleterious effect on cognition. The *FMR1-217* RNA, which is likely translated as it is associated with ribosome footprints, encodes a 31 amino acid polypeptide composed of 19 residues from exon 1 and 12 resides from the pseudo-exon. Exon 1 encodes a Tudor or agenet domain; a platform for protein-protein interactions including methylated proteins such as FUS, DDX5 and others (50–52). It is possible that this peptide acts as a competitive inhibitor of certain protein-protein interactions, thereby producing a toxic effect in the brain.

### *FMR1-217*, R-loops, and transcription elongation

We compared expression levels of RNAs from FXS WBCs where the *FMR1-217* levels varied by ∼10 fold but where the amount of *FMR1* was similar. This analysis reduced potential confounding effects of full-length *FMR1* and FMRP. One RNA that was up-regulated with high *FMR1-217* is *PIF1*, which encodes a topoisomerase that suppresses R-loops (35) and G-quadruplex formation (53). *PIF1* was also upregulated in 2 of 3 cell lines that contain *FMR1-217* while PIF1 protein was reduced in FXS cells. This disconnection between RNA and protein levels is consistent with previous reports where ribosome profiling and RNA-seq/SILAC proteomic comparisons showed widespread post-transcriptional regulation in FXS cells (37,39,54). Most importantly, the reduction of PIF1 in FXS cells is strongly correlated with *FMR1-217* R-loop formation.

R-loops are associated with nucleotide expansion diseases including FXS (19,55), and an R-loop in *FMR1* has been reported to map near the 5’ end of the *FMR1-217* pseudo-exon (56). Our data show an R-loop extends at least 200 bases into the pseudo-exon. However, a splice-switching ASO that reduces *FMR1-217* and elevates FMRP has no effect on R-loop formation. Thus, although an R-loop is closely correlated with *FMR1-217* formation, reducing *FMR1-217* does not require changes in R-loop. This result suggests that ASOs may act through a mechanism that does not involve an R-loop, for example, by altering RNA secondary structure, blocking recognition of the pseudo-exon by the splicing machinery, or by interfering with binding of a regulatory RNA-binding protein.

FXS cells showed significantly higher levels of the intron 1-pseudo-exon amplicon relative to control. This localized effect is consistent with two features of this region. First, the CGG repeat expansion promotes RNA secondary structures, including RNA G-quadruplexes, hairpins, and R-loops, which may locally impede RNA Pol II elongation (19,57–59). These structures also exclude nucleosomes and open chromatin that can further facilitate Pol II pausing at the promoter (60). Second, the pseudo-exon is flanked by a weak, non-canonical splice site rather than a strong canonical one. Slowed elongation through the repeat, combined with inefficient recognition of this weak site, would be expected to selectively accumulate unspliced pre-mRNA at the intron 1-pseudo-exon junction relative to amplicons at canonical, efficiently spliced junctions.

Total *FMR1* transcripts amplified by primers spanning the TSS region were higher in FXS compared to control cells, a finding that could reflect either increased transcription initiation of *FMR1-217* or an upstream alteration in start site usage. CGG repeat expansion has also been reported to shift *FMR1* transcription toward an upstream start site among the three annotated *FMR1* TSSs (61). Because our primers detect transcripts from all three sites, this altered TSS usage could contribute to the elevated pre-mRNA level. An additional observation was that *FMR1-217* pre-mRNA levels were already elevated in FXS cells compared to control at the 0-minute timepoint. This suggests that a subset of Pol II remains resistant to run-off and persists at the 5’ end of *FMR1*, including the pseudo-exon in FXS. Slow transcription elongation can influence alternative splicing and contribute to mis-splicing by allowing the recognition of weak splice sites. Indeed, application of camptothecin, which slows elongation by forming a physical roadblock to Pol II transit, promotes *FMR1-217* formation. Large pathogenic repeat expansions in general slow transcription and alter in splicing patterns in a number of genes (62–64). In *FMR1*, the CGG expansion forms an R-loop/G-quadruplex structure (65), which likely impedes Pol II elongation thereby causing recognition of the weak splice in the *FMR1-217* pseudo-exon.

### Splicing factors and *FMR1-217* formation

Finally, we find that PTBP1 and its paralog PTBP2 are key regulators of *FMR1-217* mis-splicing. RBPmap analysis identified predicted bindings sites in or near the *FMR1-217* pseudo-exon and CLIP-seq data further identified PTBP1 and PTBP2 binding sites within *FMR1*. Interestingly, PTBP1 and PTBP2 each show cell type- and stage-dependent effects on *FMR1-217* splicing. PTBP1/PTBP2 acts as either a repressor or activator of mis-splicing (66). Typically, PTBP1 inhibits splicing by binding its recognition site (UCUUC) and interfering with U2AF65 recognition of splice sites. It can act as an activator through binding to an internal splicing enhancer (which we have been unable to identify computationally via RBPmap), competition with other splicing factors, or other contexts (66–68).

In contrast, TDP-43 knockdown reduced both full-length *FMR1* and *FMR1-217* across the cell types tested, despite a predicted binding motif near the pseudo-exon. This indicates that TDP-43 does not specifically regulate *FMR1-217* splicing but instead contributes to general *FMR1* production. Together, these findings identify PTBP1/PTBP2 as specific regulators of *FMR1-217* pseudo-exon inclusion.

## Acknowledgements

We thank Dr. Megan Orzalli (University of Massachusetts Chan Medical School) for providing control and FXTAS human foreskin fibroblasts, Dr. Elizabeth Berry-Kravis (Rush University Medical Center) for providing FXS patient-derived fibroblasts, Dr. Sandra Almeida (University of Massachusetts Chan Medical School) for providing control iPSCs, and Dr. Peter Todd (University of Michigan) for providing FXS iPSCs. We thank Dr. Sneha Shah, Dr. Mariya Ivshina, Daniel Barnes, Dr. Ozkan Aydemir (University of Massachusetts Chan Medical School) and Dr. Juan Valcarcel (Center for Genomic Regulation, Barcelona) for valuable discussions and suggestions. The graphical abstract and illustrations were created with BioRender.com. This work was funded by the National Institutes of Health (NIH) grants R35GM149216 (NIGMS), R01NS132935 (NINDS) and the FRAXA Research Foundation to JDR.

