## Supplemental Figures and Tables for "Dysregulation of *FMR1* Splicing in Human Fragile X Syndrome"

### Figure S1

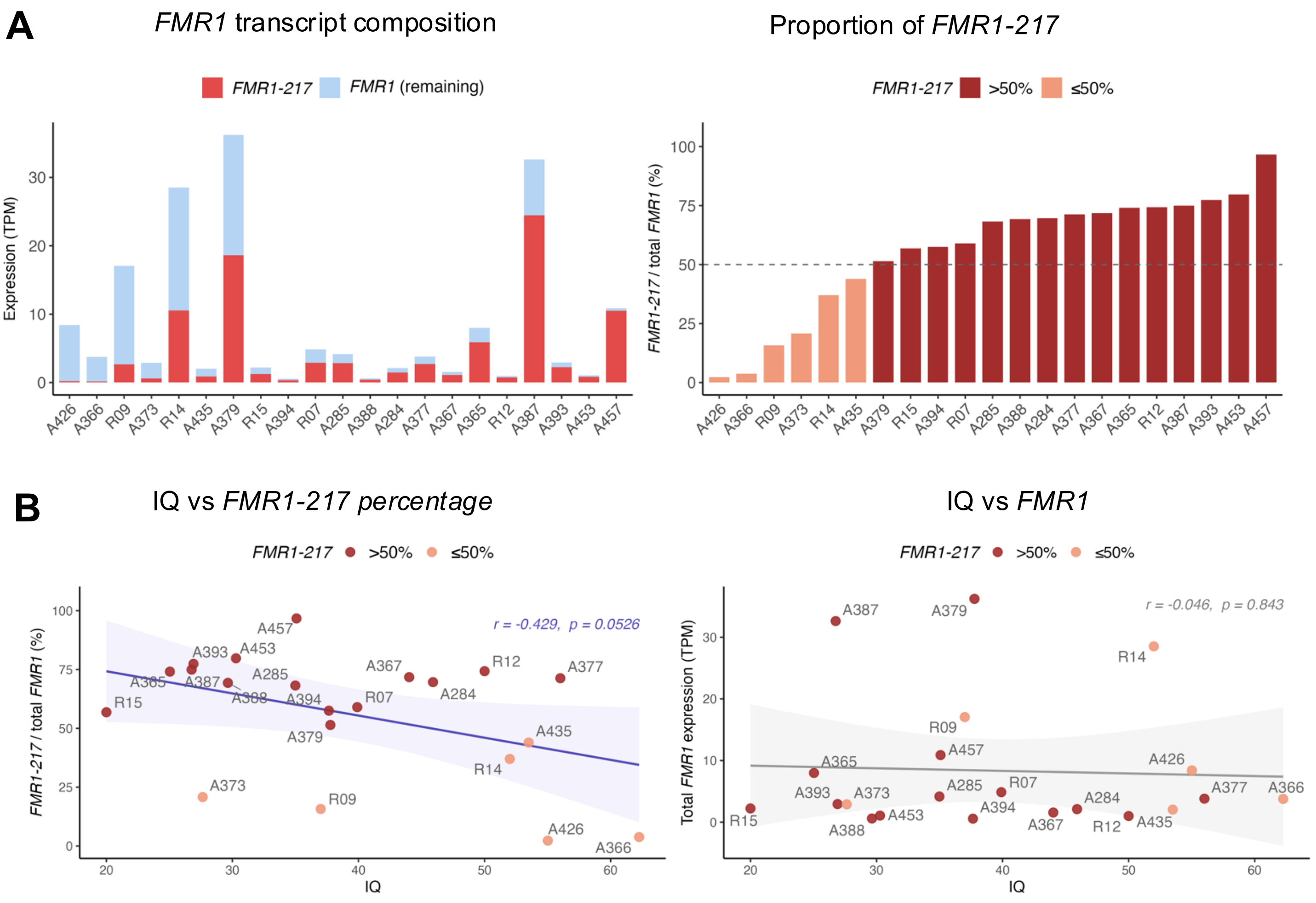


**Figure S1. Percentage of *FMR1-217* and IQ in FXS.**

(A) Transcript expression (TPM) of *FMR1-217* (dark red) and remaining *FMR1* (blue) per individual (left), and percentage of *FMR1-217* per individual (right) derived from published blood RNA-seq data (Shah et al., 2023, n = 21). Samples are ordered from lowest to highest *FMR1-217* percentage (left to right). Dashed line indicates the 50% threshold. Samples with *FMR1-217*> 50% of total are shown in dark red. Samples with FMR1-217 <= 50% are shown in salmon.

(B) Correlation between IQ and *FMR1-217* percentage (left) and total *FMR1* expression (right) in FXS individuals with detectable *FMR1* expression (n = 21). Color indicates *FMR1-217* percentage as in (A). Regression lines with 95% confidence intervals are shown. Pearson correlation coefficients and p-values are indicated. Individuals with undetectable *FMR1* were excluded from this analysis to avoid confounding by complete transcriptional silencing.

### Figure S2

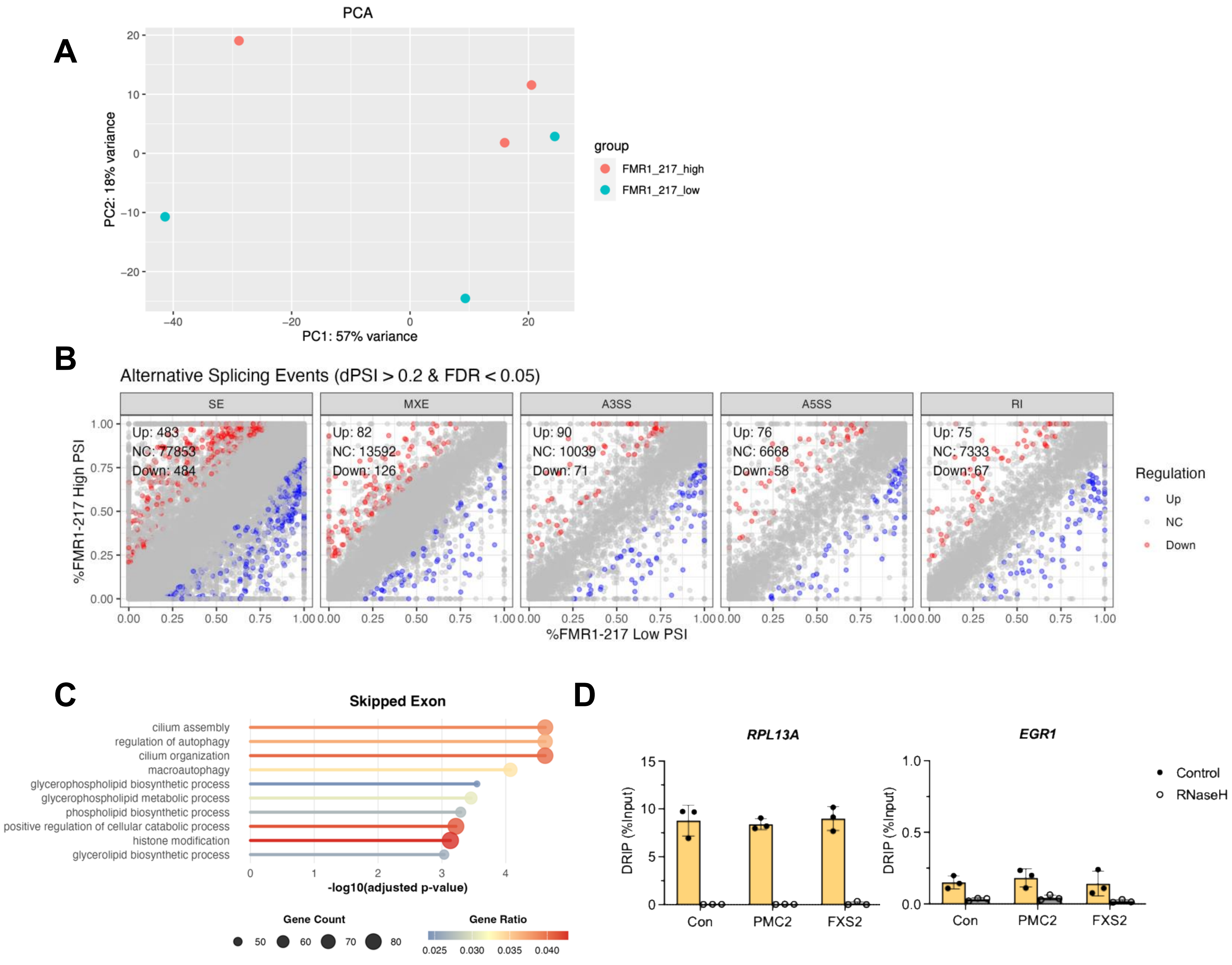


**Figure S2. Principle component and alternative splicing analysis of FXS LCLs**

(A) Principal component analysis (PCA) of RNA-seq data from *FMR1-217*-low and *FMR1-217*-high FXS LCLs.

(B) Alternative splicing analysis performed using rMATS. Five types of alternative splicing events were analyzed: skipped exons (SE), mutually exclusive exons (MXE), alternative 5′ splice sites (A5SS), alternative 3′ splice sites (A3SS), and retained introns (RI). Statistically significant inclusion events are shown in blue, and exclusion events are shown in red. Events were defined using an FDR cutoff of <0.05 and an absolute inclusion level difference (delta percent splice in, ΔPSI) >0.2.

(C) GO-term enrichment analysis of the skipped exon events.

(D) DRIP-qPCR analysis of R-loop enrichment at *RPL13A* and *EGR1* in control, premutation carrier, and FXS LCLs, with or without RNase H treatment. Error bars represent SD (n = 3).

#### Figure S3

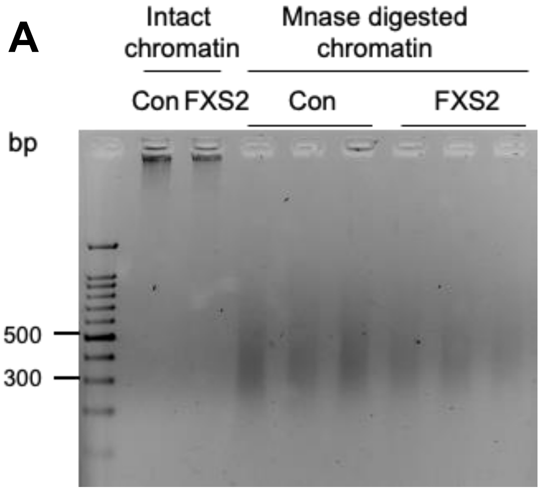


**Figure S3. Agarose gel analysis of MNase-digested chromatin**

(A) Agarose gel image showing intact chromatin and MNase-digested chromatin. Crosslinked chromatin samples were treated with 200U MNase, followed by de-crosslinking prior to gel electrophoresis. MNase digestion produced DNA fragments with predominant size range of approximately 300-500bp.

### Figure S4

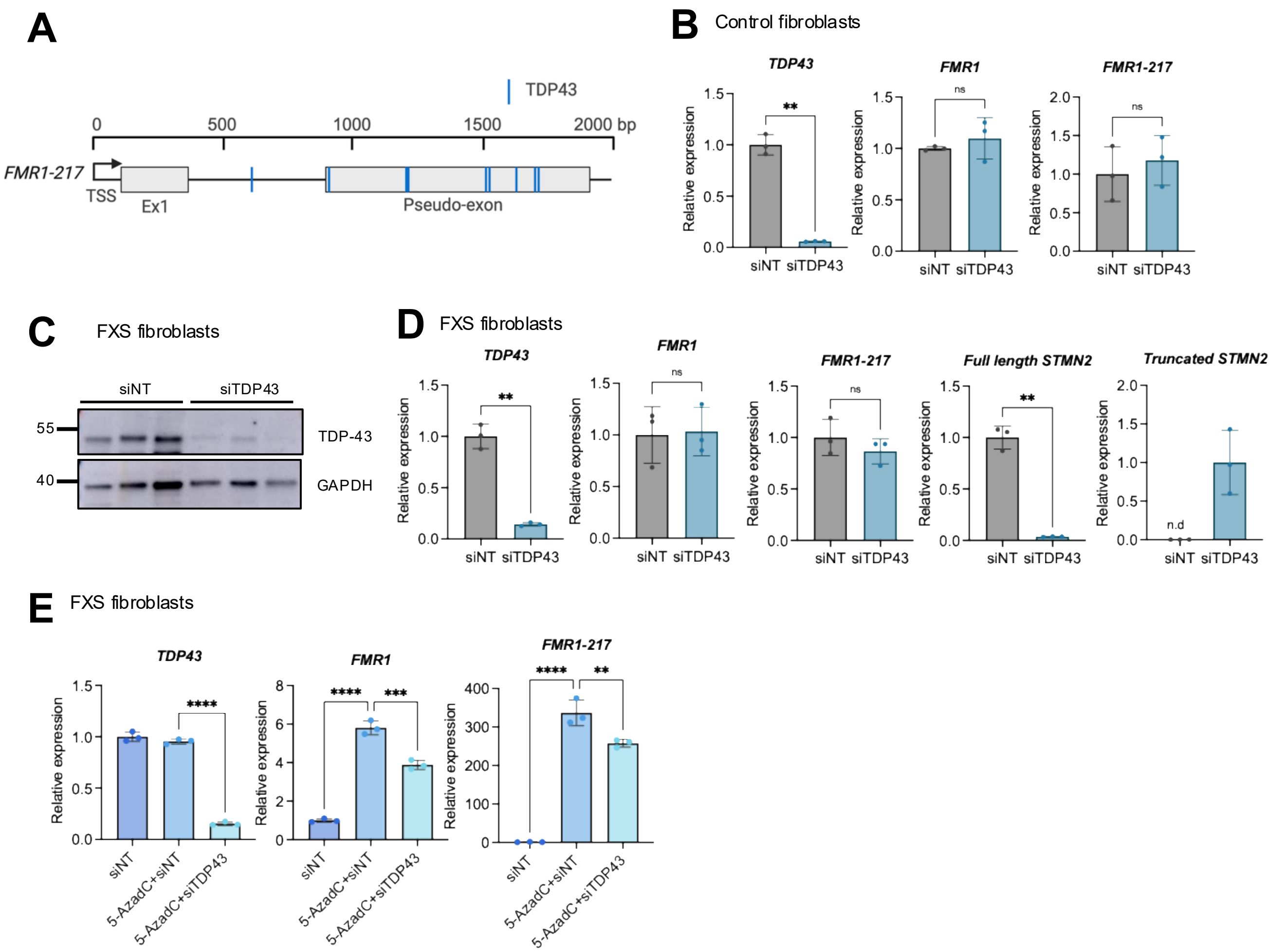


**Figure S4. TDP-43 regulates *FMR1* and *FMR1-217* expression in FXS fibroblasts.**

(A) Schematic of predicted TDP-43 binding motifs within FMR1-217.

(B) mRNA levels of *TDP-43*, *FMR1*, and *FMR1-217* in Control fibroblasts treated with non-targeting siRNA (siNT) or siTDP-43. Error bars represent SD (n = 3). Statistical significance was determined by two-tailed Student's t-test. **P < 0.01.

(C) Immunoblot showing TDP-43 knockdown in FXS fibroblasts. GAPDH was used as a loading control.

(D) mRNA levels of *TDP-43*, *FMR1*, and *FMR1-217* in FXS fibroblasts after TDP-43 knockdown. Full-length *STMN2* and truncated *STMN2* mRNAs were measured as positive controls. Error bars represent SD (n = 3). Statistical significance was determined by two-tailed Student's t-test. **P < 0.01.

(E) FXS fibroblasts were treated with 5-AzadC for 7 days and siRNAs for 3 days. mRNA levels of *TDP-43*, *FMR*1, and *FMR1-217* are shown. Error bars represent SD (n = 3). Statistical significance was determined by one-way ANOVA. **P < 0.01, ***P < 0.001, ****P < 0.0001.

#### Figure S5

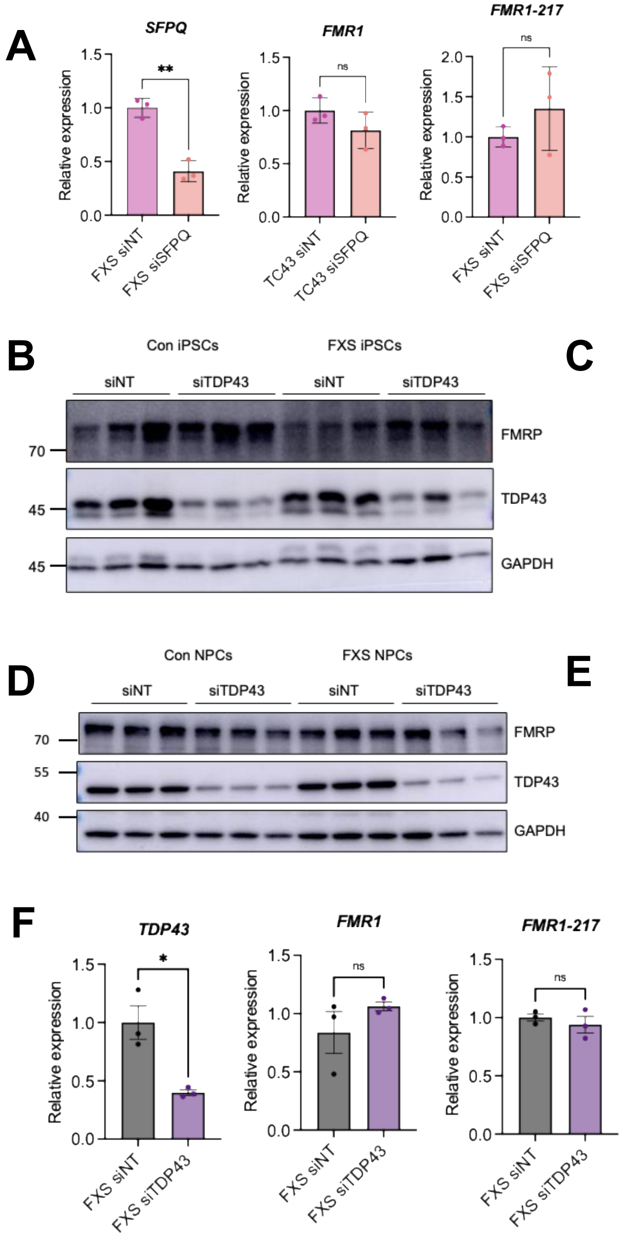


**Figure S5. TDP-43 regulates general FMR1 transcription in FXS iPSCs.**

(A) Relative mRNA levels of *SFPQ, FMR1, and FMR1-217* in FXS iPSCs treated with siNT or siSFPQ. Error bars represent SD (n = 3). Statistical significance was determined by two-tailed Student's t test. **P < 0.01.

(B) Immunoblot analysis of FMRP and TDP-43 in control and FXS iPSCs following transfection with siNT or siTARDBP. GAPDH was used as a loading control.

(C) Relative mRNA levels of *TDP-43*, *FMR1*, and *FMR1-217* in control and FXS iPSCs treated with siNT or siTARDBP. Error bars represent SD (n = 3). Statistical significance was determined by two-tailed Student's t-test. **P < 0.01, ***P < 0.001.

(D) Immunoblot analysis of FMRP, TDP-43, and GAPDH in control and FXS NPCs following transfection with siNT or siTARDBP.

(E) Relative mRNA levels of *TDP-43*, *FMR1*, and *FMR1-217* in control and FXS NPCs treated with siNT or siTARDBP. Error bars represent SD (n = 3). Statistical significance was determined by one-way ANOVA. *P < 0.05, ****P < 0.0001.

(F) Relative mRNA levels of *TDP-43*, *FMR1*, and *FMR1-217* in FXS neurons treated with siNT or siTARDBP. Error bars represent SD (n = 3). Statistical significance was determined by two-tailed Student's t-test. *P < 0.05

#### Supplementary Table

##### Table S1. Primers used for qPCR

| Gene | Primer | Sequence (5′-3′) | Reference |
|---|---|---|---|
| EGR1 | Forward | GAACGTTCAGCCTCGTTCTC | Sanz et al. (18) |
| | Reverse | GGAAGGTGGAAGGAAACACA | Sanz et al. (18) |
| FMR1 | Forward | TAGCAGGGCTGAAGAGAA | This study |
| | Reverse | TTCATGAACATCCTTTACAAATGC | This study |
| FMR1-217 | Forward | TAGCAGGGCTGAAGAGAA | This study |
| | Reverse | CAGTGGAGCTCTCCGAAGTC | This study |
| FMR1 217A | Forward | AGTAAGAAGCGGTAGTCGGC | This study |
| | Reverse | CTGGGCCATGTTAGGGTCTT | This study |
| FMR1 217B | Forward | TAAATTCAGGAATGCACATGC | Groh et al. (19) |
| | Reverse | CCTGAAGTTTCATGGCATATATT | Groh et al. (19) |
| FMR1 217C | Forward | AAACTGTTCCATACTTTGAGCAC | This study |
| | Reverse | AAACGCGGGGTTACCTTTTG | This study |
| FMR1 217D | Forward | TGGCAATAGAAGGTGCGTGT | This study |
| | Reverse | CCTATAGCCAAACGTGTCCTGT | This study |
| FMR1 Exon 1-Intron 1 | Forward | AGCCCACCTCTCGGGGG | This study |
| | Reverse | GCCCTAGAGCCAAGTACCTTGTA | This study |
| FMR1 Exon 2-Intron 2 | Forward | ACAGTTGCATTTGAAAACAAGTAAG | This study |
| | Reverse | AAGCACTCAAACTGGACTTGA | This study |
| FMR1 Intron 1-Pseudo-exon | Forward | GCCTGTCGTGTGGGTAGTTG | This study |
| | Reverse | GGAGCTCTCCGAAGTCCCAA | This study |
| FMR1 Intron 8-Exon 8 | Forward | ACATGGCTGGCCTAAATACAGT | This study |
| | Reverse | TGAAATCTCGAGGCAAGCTG | This study |
| FMR1 TSS | Forward | GAACAGCGTTGATCACGTGA | Lee et al. (20) |
| | Reverse | ACCGGAAGTGAAACCGA AAC | Lee et al. (20) |
| GAPDH promoter | Forward | TACTAGCGGTTTTACGGGCG | Yamakawa et al. (21) |
| | Reverse | TCGAACAGGAGGAGCAGAGAGCGA | Yamakawa et al. (21) |
| HPRT | Forward | CCTGGCGTCGTGATTAGTGA | This study |
| | Reverse | CGAGCAAGACGTTCAGTCCT | This study |
| PIF1 | Forward | CAATCCTCTGAGCCTCACCC | This study |
| | Reverse | CCAGGAGGTTGAGAGTCCCT | This study |
| PTBP2 | Forward | GGTAATTCCCCATTGCATCGT | This study |
| | Reverse | GATAGGTGAAGGGTGGCAGA | This study |
| RPL13A | Forward | AGGTGCCTTGCTCACAGAGT | Sanz et al. (18) |
| | Reverse | GGTTGCATTGCCCTCATTAC | Sanz et al. (18) |
| STMN2 (Full-length) | Forward | AGCTGTCCATGCTGTCACTG | Baughn et al. (22) |
| | Reverse | GGTGGCTTCAAGATCAGCTC | Baughn et al. (22) |
| STMN2 (Truncated) | Forward | CTTTCTCTAGCACGGTCCCAC | Baughn et al. (22) |
| | Reverse | ATGCTCACACAGAGAGCCAAATTC | Baughn et al. (22) |
| TARDBP | Forward | GTGGGCTTCGCTACAGGAAT | This study |
| | Reverse | GATTTCCCCAGCCAGCATCT | This study |

##### Table S2. Primers used for FMR1-217-3xFLAG cloning

| Gene | Sequence (5′-3′) | Reference |
|---|---|---|
| 217F | CCGGAATTCGCCACCATGGAGGAGCTGGTGGTGGAAG | This study |
| 217R | CGCCACCCAGCCCTCGCCCAGAACAGTGGAG | This study |
| 3XFLAG-R | CGCGGAGATCTTTACTTGTCGTCATCGTCTTTGTAGTCACCCTTGTCGT CATCGTCTTTGTAGTCACCCTTGTCGTCATCGTCTTTGTAGTCGGAGCC GCCGCCACCCAGCCCTCGCCCAGAACAGTGGAG | This study |
